# Common Permutation Methods in Animal Social Network Analysis Do Not Control for Non-independence

**DOI:** 10.1101/2021.06.04.447124

**Authors:** Jordan D. A. Hart, Michael N. Weiss, Lauren J. N. Brent, Daniel W. Franks

**Affiliations:** Centre for Research in Animal Behaviour, University of Exeter, UK; Center for Whale Research, Friday Harbour, WA, USA; Departments of Biology and Computer Science, University of York, UK

## Abstract

The non-independence of social network data is a cause for concern among behavioural ecologists conducting social network analysis. This has led to the adoption of several permutation-based methods for testing common hypotheses. One of the most common types of analysis is nodal regression, where the relationships between node-level network metrics and nodal covariates are analysed using a permutation technique known as node-label permutations. We show that, contrary to accepted wisdom, node-label permutations do not automatically account for the non-independences assumed to exist in network data, because regression-based permutation tests still assume exchangeability of residuals. The same assumption also applies to the quadratic assignment procedure (QAP), a permutation-based method often used for conducting dyadic regression. We highlight that node-label permutations produce the same *p*-values as equivalent parametric regression models, but that in the presence of non-independence, parametric regression models can also produce accurate effect size estimates. We also note that QAP only controls for a specific type of non-independence between edges that are connected to the same nodes, and that appropriate parametric regression models are also able to account for this type of non-independence. Based on this, we suggest that standard parametric models could be used in the place of permutation-based methods. Moving away from permutation-based methods could have several benefits, including reducing over-reliance on *p*-values, generating more reliable effect size estimates, and facilitating the adoption of alternative types of statistical analysis.

## Introduction

Social network analysis is a central tool in the study of animal sociality. Social networks characterise the structure of social connections between individuals, and are useful for answering a wide range of biological questions related to social structure, the evolution of sociality, information and disease transmission, and more (Farine and Whitehead, 2015). Social networks are usually analysed quantitatively at three levels: global, nodal, or dyadic. Global network metrics characterise features of the entire network, such as connection density or longest path; nodal metrics describe each node’s position in the network relative to the other nodes; and dyadic metrics describe each edge’s position in the network (Butts, 2008). Two common types of hypotheses in animal social network analysis can be characterised as: ‘nodal metrics are related to nodal covariates’, and ‘the presence or metric of edges are related to dyadic covariates’ (Croft et al., 2011; Dekker et al., 2007). The types of analyses used to test these hypotheses are known by various names, but we will refer to them as *nodal regression* and *dyadic regression* respectively. These analyses usually use permutation-based regression techniques such as node-label permutations or the quadratic assignment procedure (QAP). Node-label permutations have typically been applied to nodal regression and QAP to dyadic regression (Farine, 2017). The justification for the use of permutation-based regression tests over parametric regression models is that network data are inherently non-independent and therefore break the assumptions of parametric regression.

### The problem of non-independence

Many conventional statistical analyses make the assumption that data are independent (Cohen, 1992). This assumption is key to reliable data analysis because it defines the source and nature of noise in data generating processes, and is therefore closely linked to null hypothesis significance testing and calculation of *p*-values. In the case of regression analysis, a noise term is included in the model to account for non-systematic, independent random noise present in the data (Draper and Smith, 1998). This assumption is convenient because it has appealing mathematical properties, but in practice can rarely be met. In the presence of known sources of non-independence, statisticians often use explicit models of the sources of non-independence, for example autocorrelation models are frequently deployed in time series analysis to account for the known temporal dependencies in sequential data (Wei, 2013).

In network data, dependencies are assumed to be more complex. A common example is that undirected node strength is explicitly related to the node strength of every other node in the network, even for nodes that are not directly connected to the node of interest (Sosa et al., 2021). Therefore noise in the data may be linked to various structural features of the network and would be poorly modelled by an independent noise term. Whether or not the *p*-value of a statistical analysis can be trusted depends on how well the process that generates noise in the data is described by the model, which in the case of parametric regression models requires independent residuals.

Inappropriate noise terms in statistical models are a major problem when scientific hypotheses are evaluated using null hypothesis significance testing (Anderson and Robinson, 2001). Null hypothesis significance testing is based on the concept of constructing a null model that describes the data if there is no relationship between variables of interest, and that any relationship between them is due to chance alone (Wasserman, 2004). Tests are usually conducted by calculating the *p*-value, which is the probability of getting coefficient estimates at least as extreme as those from hypothetical data generated under the null hypothesis. Parametric regression tests use the noise term in the model to estimate what coefficient values are likely ‘by chance’, and to subsequently calculate the *p*-value. If the noise term in the regression model does not approximately match the process that generates noise in the data generating process, then the *p*-value will not reflect what is expected by chance, and therefore will not be reliable.

### Permutation tests

The interconnectedness of social networks appears to break the independence assumptions of parametric regression models. This has been a long term concern of behavioural ecologists conducting social network analyses (Croft et al., 2010). Because permutation tests relax some assumptions about the distributions of noise terms, they have been widely adopted with the aim of enabling regression analysis in the presence of non-independence (Croft et al., 2011). The notion behind permutation regression tests is that if there is no effect, nodal (or dyadic) covariates are equally likely to belong to any node (or dyad). When using node-label permutations, parametric regression is applied to the network and a test statistic such as the coefficient estimate or *t*-value is recorded. Then the node labels are swapped at random and the test statistic is re-estimated from the new dataset with permuted node labels. This permutation step is repeated many times to build a distribution of test statistic values under the null hypothesis of no relationship between node centrality and nodal covariate. The observed test statistic can then be compared to the null distribution to calculate statistical significance.

In permutation tests, some confounds can be accounted for by constraining permutations to between certain data points (Winkler et al., 2015). Constraining permutations is the key notion behind QAP, which works in much the same way as node-label permutations, but because the dyad is the unit of analysis, relabelling nodes effectively permutes all connections of a node at the same time. This controls for any dependence between edges that connect to the same node. Constraining permutations in this way means that the model that calculates the observed test statistic does not account for the confounds being used as constraints, and subsequently does not take them into account when calculating effect size estimates. Consequently, effect size estimates computed in this way will be incorrect, to the extent that they may even have the wrong sign (Franks et al., 2021).

Instead of explicitly assuming a parametric noise term, permutation tests assume that under the null hypothesis, any rearrangement of the data is equally likely (Good, 2000). In a regression setting this generates the null hypothesis of no relationship between the response and covariates. Thus, permutations have the benefit of removing the need for some assumptions about the distributions of noise in data generating processes. The assumption that all permutations of the data must be equally likely under the null hypothesis is known as exchangeability of data points. This means that data points must be freely exchangeable under the null hypothesis without changing their joint probability, which depends on the underlying dependence structure of the data points. In the presence of dependence between data points, unconstrained permutations of the data do not preserve dependence structure (see Figure 1). This breaks the exchangeability assumption of permutation tests for much the same reason as non-independence breaks the assumptions of parametric regression (Winkler et al., 2015). This is illustrated in Figure 1a where the data points 1 and 2 are independent, but data points 1 and 3, and 2 and 3 are dependent on each other. This forms a dependence structure that must not be broken by permutations, but node-label permutations freely permute data points and thus break any dependence structure in the data.

**Figure 1:**
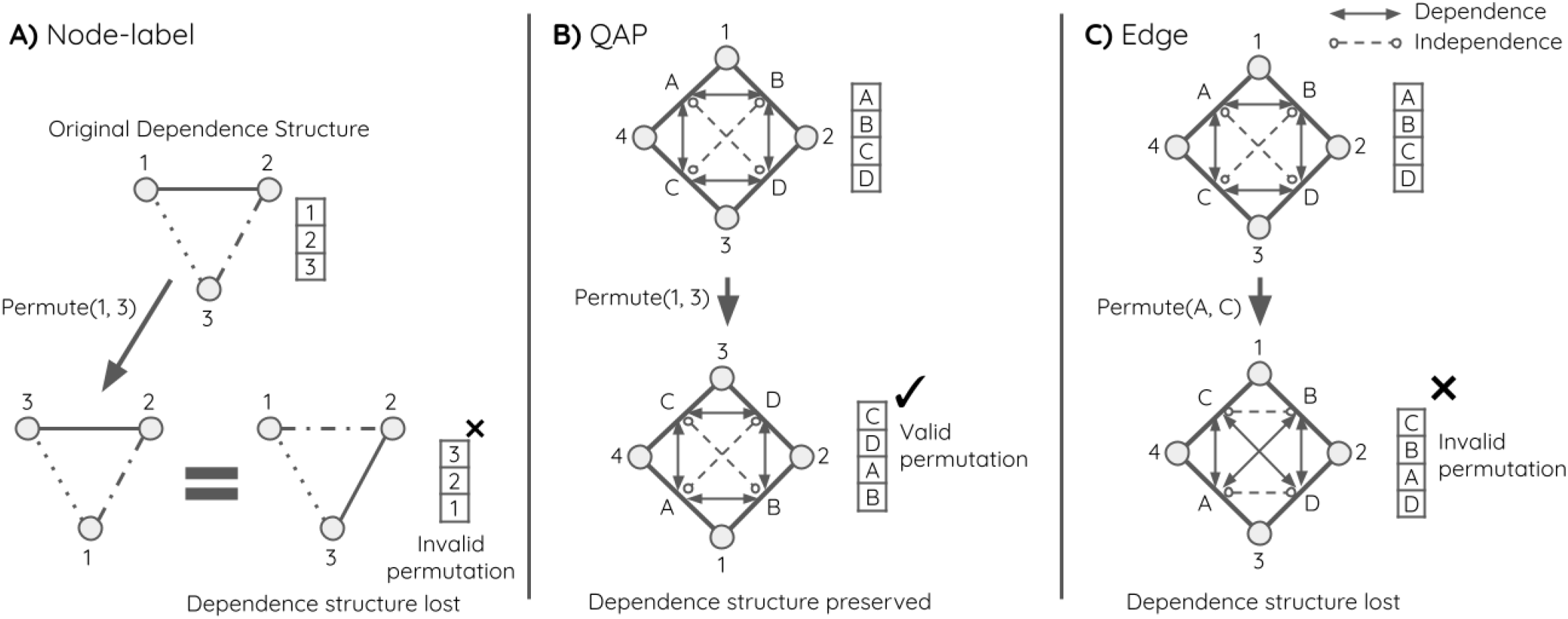
Dependence structure between data points must not change under permutations. In node-label permutations (A), any dependency structure between nodes would be lost when permuting. This could break the exchangeability assumption and generate invalid permutations in the presence of a strong dependence structure. In QAP (B), node labels are again permuted, but at a dyadic level this is equivalent to permuting multiple edges at once, which preserves the assumed dependency structure of the data, generating valid permutations and correct p-values. Hypothetically if edges were permuted freely (C), the pattern of the dependency structure in the original data would be lost, and the resulting permutations of the data would not be valid. Solid lines with arrows on both ends denote dependence between data points, and dashed lines with circles on both ends denote independence between data points. Note that some dependencies have been omitted for clarity.

The exchangeability condition also applies to QAP, though QAP makes the explicit assumption that dyads are dependent on the nodes to which they are attached. This assumption means that the QAP controls for one specific type of non-independence, but is not immune from more complex dependencies such as dyads depending on other aspects of network substructure. Figure 1b shows how QAP restricts permutations on networks to move multiple edges at once, preserving the original dependency structure. Hypothetically speaking, if in QAP edges were permuted freely, as nodes are in node-label permutations, the dependency structure would not be preserved, and invalid permutations would be generated (as shown in Figure 1c). Therefore permutation tests do not automatically correct for non-independence, meaning node-label permutations will produce equivalent p-values to comparable parametric regressions, and QAP will provide equivalent p-values to comparable to parametric regressions with a term for node dependence (Good, 2000).

In this paper we provide examples to illustrate that, in practice, node-label permutations and parametric regression yield the same true and false positive rates. We show that QAP correctly accounts for a specific type of non-independence, but that alternative non-permutation models are also capable of accounting for such non-independence. We also show that in the presence of non-independence that is not explicitly accounted for, both node-label permutations and QAP yield inflated false positive rates, highlighting that permutations do not automatically control for non-independence. Finally we discuss the potential benefits of using standard parametric models for regression analysis on network data.

## Methods

In this section, we use network simulations to illustrate that node-label permutations achieve the same true positive rates (power) and false positive rates (type I error) as ordinary least squares in nodal regression to detect trait-based differences in a common node-based measure of centrality. We also use simulations to show that network substructure can introduce dependence structure in the data that neither node-label permutations nor QAP can account for. Finally, to demonstrate that parametric statistical models are able to account for specific types of non-independence in the same way as QAP, we compare QAP to both ordinary least squares and a multimembership linear model that includes a node dependence term.

### Simulations: Nodal regression

#### Trait-based strength differences

To demonstrate that node-label permutations perform the same as parametric regression, we compared a standard OLS model (LM) to node-label permutations where OLS is used to calculate the test statistics. To generate the data, we used the simulation model described by Farine and Whitehead (2015). The simulations assigned a gregariousness score to each individual in the population of size *n* = 20 from a Poisson distribution. Individuals were then assigned a sex either according to their gregariousness (effect), or at random (no effect). Sampling periods were simulated where the probability of a pair interacting in a sampling period was proportional to the combined gregariousness scores of the two individuals, giving a weighted, undirected network. Node strength was calculated as the sum of each node’s connection strengths. Node strength was regressed against sex using OLS. Node-label permutations were conducted with 10,000 permutations on the networks to generate the null distribution using the slope coefficient as the test statistic. The observed coefficient was compared to the null distribution to compute a two-sided *p*-value and effect size estimate for the null hypothesis of no effect. This was repeated 1,000 times in the presence of both an effect and no effect, and the true positive and false positive rates were computed.

#### Nodal dependence on clique membership

The non-independence of network data can take many forms, but to demonstrate one possible form, we considered the case where a network is formed from two unknown underlying cliques. Our simulations assigned nodes to one of two cliques at random, with equal probability of being assigned to either clique. Dyads of nodes that were in the same cliques had an 80% chance of having a non-zero edge, whereas dyads of nodes that were in different cliques only had a 40% chance of having a non-zero edge. Edge weights were drawn from a uniform U(0, 1) distribution. Nodal covariates were assigned according to a linear combination of node strength, a clique dependence variable, and a random noise term, drawn from a uniform distribution U(0, 1). The clique dependence variables were drawn from a uniform distribution U(0, 1), and were used to create an effect of clique membership on nodal covariates. If no effect was being simulated, the coefficient of node strength was set to zero to remove the effect, otherwise it was set to 0.05. This simulation creates an effect of non-independence because under null hypothesis, the size of the cliques will affect node strength, and clique membership affects the nodal covariates. The strength of a node will depend on the size of its clique, which is generated by a stochastic process, so there is potential for spurious correlation between node strengths and nodal covariates. This simulation is designed to simulate the effect of substructures in the network that may be difficult or even impossible to detect either manually or computationally. The simulation was repeated 1,000 times with and without the effect, and the two-sided *p*-values and effect size estimates for each method were recorded.

### Simulations: Dyadic regression

#### Dyadic dependence on nodes

To demonstrate the performance of QAP against parametric regression, we designed simulations based on those described by Dekker et al. (2007). Simulations were carried out by simulating the response and predictor matrices as being partially dependent on a node-level vector:

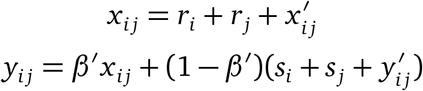

where *x* and *y* are the observed matrices, *r* and *s* are the node dependencies for *x* and *y* respectively, *y*^*′*^ and *x*^*′*^ are the true, underlying social preferences, and *β*^*′*^ describes the relationship between *x* and *y*. This creates a relationship between *x* and *y* when *β*^*′*^ ≠ 0. The matrices were symmetric of size *n* = 20, with elements drawn from a uniform *U* (0, 1) distribution. The node dependence vectors *r* and *s* were also drawn from a uniform *U* (0, 1) distribution, and the effect parameter *β*^*′*^ was set to either *β*^*′*^ = 0 to simulate no effect, or to *β*^*′*^ = 0.20 to simulate a moderate effect. In line with Dekker et al. (2007), intercepts were not included in the simulation or model, but this does not affect the generality of the results.

Previous studies have demonstrated that QAP is effective at accounting for node dependencies in dyadic regression (Dekker et al., 2007). The reason for this is not because QAP is a permutation test, but because QAP makes explicit assumptions about the sources of non-independence. This same assumption can also be built into parametric models using random effects. Since each edge depends on two nodes, and each two nodes have only one edge between them, conventional random effects cannot be used to control for node dependence, as this would use one random effect per unit of analysis (per dyad). Instead a random effect is used for each node, and the random effects for two nodes that each edge is between are included in the model. This type of mixed model is often referred to as a multimembership model (Rushmore et al., 2013; Boyland et al., 2016). We implement the following multimembership linear model:

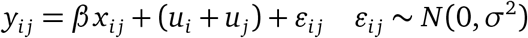

where *x, y* are the predictor and response matrices, *u* is a random effect vector describing the influence of each node on its connected dyads, and *ε* is an independent noise term. The vector *u* is treated as a set of parameters to be learned, which introduces a considerable number of parameters to the model. For computational reasons, the model was fit using numerical least squares with the optim function in R, but these types of models are also supported in R packages such as brms and MCMCglmm (R Core Team, 2020; Bürkner, 2018; Hadfield, 2010).

The simulated matrices *x* and *y* were regressed against each other using the following three methods: a standard OLS linear model (LM), QAP, and the multimembership linear model described previously (MMLM). The *p*-values and effect size estimates from each were recorded. The QAP method used 1,000 permutations to generate the null distribution. As with the previous simulations, this was repeated 1,000 times in the presence of both an effect and no effect, and true positive and false positive rates were computed.

#### Dyadic dependence on clique membership

As with the simulation of the effect of network substructure on nodal regression, the aim of this simulation was to demonstrate how dependence on network substructures can affect the performance of dyadic regression. To demonstrate the potentially subtle nature of non-independence in dyadic regression, we introduce dependence in a different way to the nodal simulation. In this simulation we assume that fully connected subgraphs of size 4 form cliques that affect both the strengths of edges and dyadic covariates for dyads in the cliques. Naturally a dyad may belong to multiple cliques, so may have a complex structure of dependencies. Cliques of size 4 are used because they are the smallest possible subgraph that does not follow the assumptions of QAP. The rest of the simulation proceeds in the same way as the previous simulation with the three models: LM, MMLM, and QAP.

## Results

Plots of the distributions of *p*-values in the presence and absence of an effect are shown for each of the four simulations in Figure 2. Under the null hypothesis of no effect the *p*-values should be uniformly distributed, whereas under the alternative hypothesis the *p*-values should be concentrated towards zero (Wasserman, 2004). The distributions of effect size estimates for the dyadic regression simulations are shown in Figure 3, and should be centered around 0.2 when there is an effect, and centered around zero when there is no effect.

**Figure 2:**
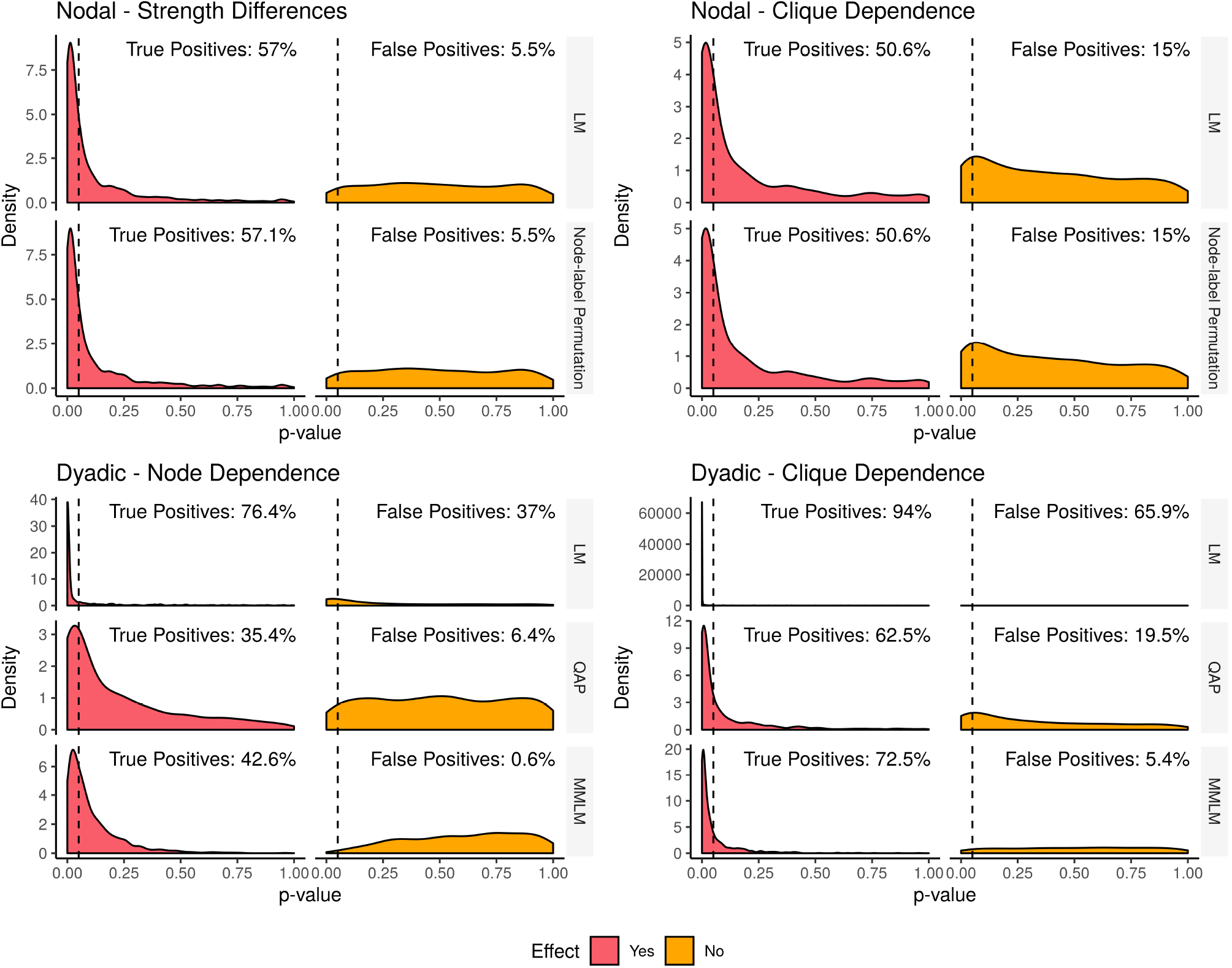
Distributions of *p*-values in nodal and dyadic regression from simulations comparing ordinary least squares regression (LM) and its permutation-based equivalents: node-label permutation and QAP respectively. The dashed line indicates the conventional significance threshold of *p* = 0.05. When there is no effect, distributions should be uniformly distributed, but when there is an effect, *p*-values should be concentrated towards zero. In the two nodal regression simulations the *p*-value distributions for both methods are identical both with and without an effect, indicating equivalent performance for both methods. Furthermore, in the presence of clique dependence in nodal regression, an inflated false positive rate is seen in the *p*-value distribution. In dyadic regression LM had a high false positive rate in both simulations, though suffered even higher false positives in the presence of clique dependence. In contrast, both QAP and MMLM performed well in the presence of node dependence. In the presence of clique dependence QAP had an inflated false positive rate of 19.5%, compared to the MMLM which had a low false positive rate of 5.4%.

**Figure 3:**
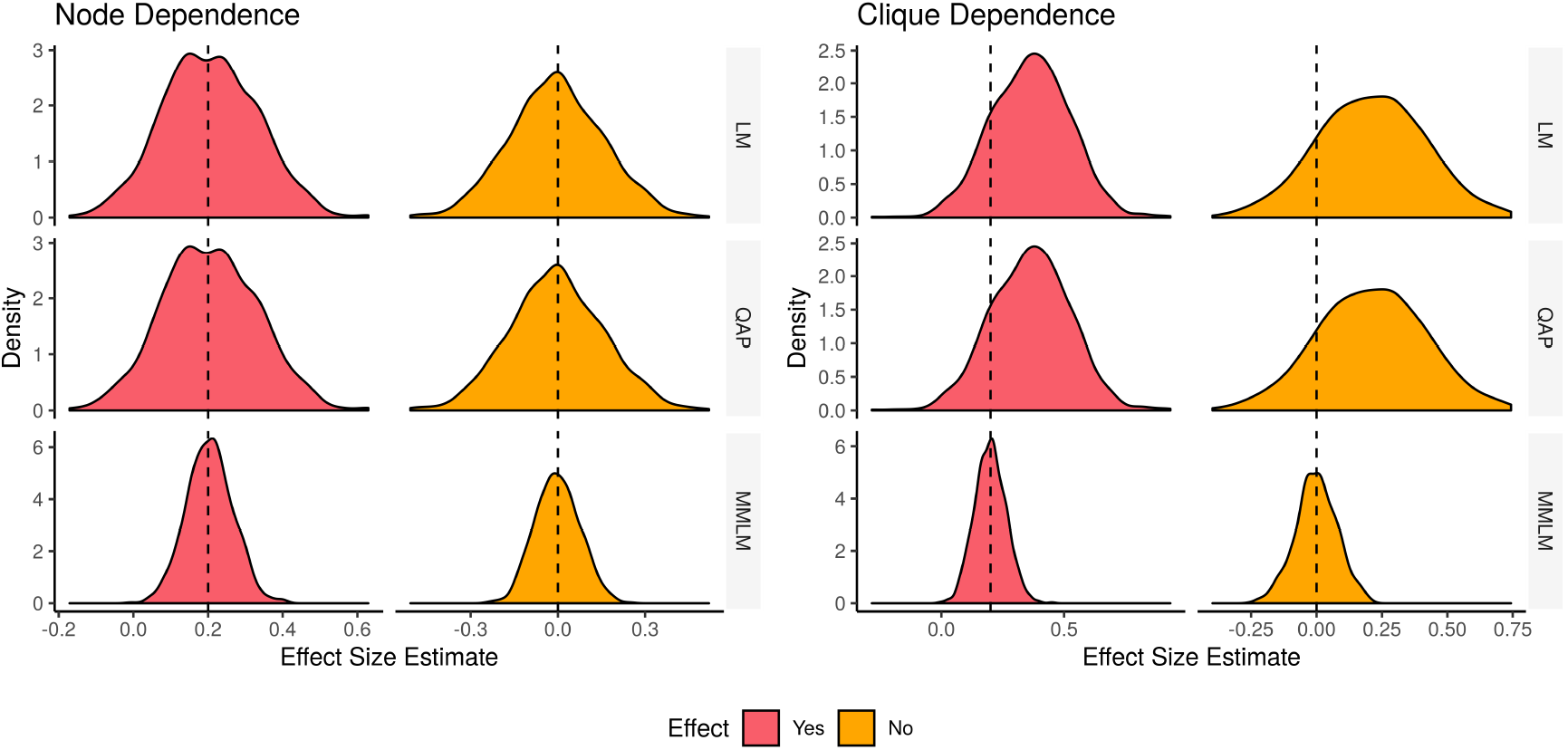
Distributions of estimated effect sizes in the two dyadic regression simulations. The LM and QAP both use the estimates directly from OLS, and therefore have the same distributions of effect sizes. The MMLM takes the dependence terms into account when making effect size estimates, and has a narrower distribution around the median effect estimates than the LM and QAP. The LM and QAP produce the same estimates, which are correct in the node dependence simulations, but biased towards a positive effect size when there is no true effect in the clique dependence simulations. The MMLM performed correctly in both scenarios and achieved a narrower distribution. The dashed line indicates the desired effect size estimate of 0.2 when there is an effect, and 0.0 when there is no effect.

### Nodal regression

#### Trait-based strength differences

In our simulations of trait-based strength differences, the LM and node-label permutation methods had true positive rates of 57.0% and 57.1% respectively, and both methods had a false positive rate of 5.5%. The distribution of *p*-values was almost identical for both methods, and under the null hypothesis was approximately uniform.

#### Nodal dependence on clique membership

When the effect was present, the two methods achieved true positive rates of 50.6% for both the LM and node-label permutations. As with the previous simulation, the LM and node-label permutations achieved the same *p*-value distributions both in the presence and absence of an effect. Unlike in the previous simulation, the *p*-value distribution was not uniform under the null hypothesis, with the methods giving inflated false positive rates of 13.7% and 13.3% for the LM and node-label permutations respectively.

### Dyadic regression

#### Dyadic dependence on nodes

The dyadic regression simulations where dyads were dependent on nodes showed that the LM method had a high true positive rate of 76.4%, QAP was more conservative with a true positive rate of 35.4%, and the MMLM had a true positive rate of 42.6%. The LM had a highly inflated false positive rate of 37.0%, compared to QAP with a false positive rate of 6.4% and the MMLM at 0.6%. QAP had approximately uniformly distributed *p*-values under the null hypothesis.

When there was an effect, the distribution of effect size estimates for the LM and QAP had a median of 0.205, with a 95% interval of (−0.0349, 0.455), compared to the effect size estimates of the MMLM, with a median of 0.200 and a 95% interval of (0.0810, 0.330). The effect size estimates from the LM and QAP had the wrong sign in 4.9% of cases, and significant results had the wrong sign in 0.4% of cases for the LM, but never for QAP. The MMLM effect size estimates had the wrong sign in 0.1% of cases, again none of these cases were statistically significant. In the absence of an effect, the distribution of effect size estimates for the LM and QAP had a median of -0.00498 with a 95% interval of (−0.315, 0.317), whereas the effect size estimates of the MMLM had a median of -0.00355 with a 95% interval of (−0.140, 0.150).

#### Dyadic dependence on clique membership

When the assumptions of QAP were broken by allowing dyads to depend on cliques in the network, in the presence of an effect, the LM achieved a true positive rate of 94.0%, compared to QAP with a true positive rate of 62.5%, and MMLM with a true positive rate of 72.5%. In the absence of an effect, the LM suffered an inflated false positive rate of 65.9%, QAP obtained a false positive rate of 19.5%, and the MMLM obtained a false positive rate of 5.4%.

In these simulations, when an effect was present, the distribution of effect size estimates of the LM and QAP had a median of 0.368, with 95% interval of (0.0453, 0.673), whereas the MMLM had a median of 0.200 with a 95% interval of (0.0790, 0.330). The LM and QAP had the wrong effect sign in 0.9% of cases. These wrong effect signs accompanied significant *p*-values in 0.2% of cases for the LM, and in zero cases for QAP. The MMLM had wrong effect signs in 0.1% of cases, and never accompanying significant *p*-values. In the case where no effect was present, the distribution of effect size estimates for the LM and QAP had a median of 0.210, with a 95% interval of (−0.196, 0.615). The distribution for the MMLM had a median of -0.000126 with a 95% interval of (−0.160, 0.160).

## Discussion

Node-label permutations and QAP are some of the most popular statistical tools used in animal social network analysis (Farine, 2017). We have highlighted that node-label permutations do not control for the non-independence of network data. We have also demonstrated that while the QAP does control for some types of non-independence, such control can also be achieved by a relatively simple parametric regression model. Additionally, we have shown that in plausible scenarios of non-independence, both node-label permutations and QAP can yield inflated levels of false positives and low statistical power, and that even properly constrained permutation models provide unreliable effect size estimates. In this section we will discuss the consequences of these findings, the potential benefits of parametric models for network analysis, and future directions for statistical analysis of networks.

### Simulation results

We illustrated that node-label permutations yielded near-identical results to the LM model on the nodal regression simulations. Node-label permutations are the non-parametric equivalent to standard regression, which are identical models when distributional assumptions are met (Good, 2000). Both methods achieved the correct false positive rates, showing that the assumptions of the models were not severely broken, and the noise term in the LM was an appropriate model for the noise. In our second nodal regression simulation we showed that in the presence of non-independence due to network substructure, the assumptions of both the LM and node-label permutations were broken, leading to inflated false positive rates. In the simulation, group size was distributed according to a random binomial process, and since group size affected network strength, this led to spurious correlation in the regression. The noise term of the LM is not designed to absorb error of this nature, and failed to produce correct *p*-values. The node-label permutation failed in the exact same way because the assumption of exchangeability of data points was broken.

Our dyadic regression simulations illustrated how the LM suffered from inflated false positive error rates because of simulated node dependence, whereas QAP and the MMLM achieved correct false positive and true positive rates. This demonstrates that while QAP does control for node dependence, the same control can be replicated by including terms for dependencies in what many researchers may consider to be more conventional statistical models. Modelling dependencies in this way is more powerful because it allows effect size estimates to fully account for non-independence, whereas QAP generates the same unadjusted effect size estimates as the LM. Higher-order nodal dependencies, or other structural dependencies are not controlled for by either QAP or any other model, unless explicitly specified. To demonstrate this, our final simulation assigned edge values and dyadic covariates according to their membership of cliques. This created a dyadic dependence on substructure that the QAP is not designed to control for. The results of the simulation agreed with the theory, showing that QAP only accounts for non-independence between adjacent dyads. The MMLM achieved a low false positive rate and a high true positive rate on this simulation, suggesting that the multimembership term was able to effectively model dyadic dependence on cliques.

### Impacts of non-independence in network analysis

The findings of this paper may raise questions about the reliability of *p*-values and effect size estimates in statistical analysis of networks. In nodal regression, where centrality metrics are regressed against nodal covariates, noise in the relationship between centrality and trait can be attributed to either measurement error in the traits or due to traits being noisy proxies for the true variables of interest. Therefore noise in the relationship can be considered to be independent between nodes, and standard regression will be an appropriate type of model for conducting nodal regression, as is widely used in several other fields (Wasserman, 2004; O’Malley and Marsden, 2008; Morselli et al., 2013; Morelli et al., 2017).

In dyadic regression, whether node dependence is a realistic assumption will again depend on the biological question and data. Where dyadic covariates are related to attributes of the nodes, such as age or sex differences, accounting for dependence on nodes will be of vital importance. This is because network structure affects both the dyadic response and dyadic covariates, creating a non-causal association between response and covariate and breaking the independence assumption. Conversely, if dyadic covariates are not dependent on nodes, the independence assumption of standard regression will hold, and QAP and multimembership models will not be necessary.

Permutation tests are also used outside of regression contexts, but exchangeability of data points is a condition of any permutation test, including datastream permutations, where raw observations of association and interactions are randomised (Bejder et al., 1998). Whether other common permutation tests are valid will depend on both the data and the biological question. For example, tests such as Bejder et al. (1998)’s test of non-random association are valid because the null hypothesis is that social associations are random, and therefore observations of individuals are assumed to not be dependent on the observations of other individuals. This means that the data points are exchangeable under the null hypothesis, and may be freely permuted.

### Future directions

The prevalence in empirical data of the types of non-independence described in our simulations is unknown. Whether or not this type of non-independence is a major issue for statistical analysis of network data will require further investigation. It is worth noting that higher-order dependencies such as clique membership may lead to apparent effects may in fact be the object of interest for many network analyses. Statistically controlling for those dependencies would remove the effect of interest, so further consideration of the role of dependencies in network data will play an important role in shaping how hypotheses can be tested in network analysis. In cases where higher-order dependencies are not the objects of interest, one potential direction for future work is to categorise potential sources of network dependence (see e.g. Tranmer et al., 2014). This would make it possible to use causal models to identify likely sources of dependence in network data and account for them statistically (Pearl, 2010). An ideal family of models for this type of analysis are mixed models, also known as multilevel, hierarchical, or random effects models, among others (Congdon, 2020). These flexible and powerful models allow for explicit modelling of interdependence between data points at multiple levels. Furthermore, other sources of noise such as uneven sampling and sampling biases could also be accounted for directly in well-specified mixed models, and would lead to more powerful, efficient, and reliable statistical analyses.

The problem of non-independence in network data has also been considered for several other types of statistical model. Exponential random graph models (ERGMs) make the same base assumption as QAP, namely that edges that are not connected to the same nodes are independent. However, extensions of ERGMs have been developed that explicitly model dependence between clique-like triangular substructures (Snijders et al., 2006; Hunter, 2007). Another example of dependence modelling in dyadic data is the actor-partner interdependence model (Cook and Kenny, 2005). The actor-partner interdependence model is a dyadic-level model that assumes that an actor’s nodal covariate depends on both the dyad and partner node. A related idea where non-independence is treated as the object of interest in analysis is the network autocorrelation model and its variants (Dittrich et al., 2020). Network autocorrelation models treat non-independence as the object of analysis rather than a nuisance factor, and have a long history of being used to test hypotheses about social influence in networks (Doreian, 1981). The common thread between these methods is that dependencies in the network are explicitly accounted for in the statistical model. We believe that approaching the problem of non-independence in network data in this way will lead to more robust analyses of animal social networks.

Over-reliance on p-values and significance testing has garnered widespread criticism over several decades (Cohen, 1994). A key drawback of p-values is that they do not indicate the magnitude or direction of an effect. For example, in some cases a significant result (p < 0.05) might be accompanied with a miniscule effect estimate that would be of little interest to the researcher. Critics of this use of the p-value have proposed to instead focus on effect size estimates and confidence intervals, and to use p-values as a complementary piece of information when drawing conclusions about analyses. Permutation tests generate effect size estimates using standard regression, so even when permutations are properly constrained to account for confounds, though they will generate correct p-values, the effect size estimates will not account for these confounds and may be unreliable (Franks et al., 2021). In our simulations the distribution of effect size estimates for QAP was around twice the width of those of the MMLM. In real world use, this could severely reduce the reliability of inference, and even introduces the possibility of statistical significance for effect sizes with the wrong sign. We suggest that future analyses could focus on accurately estimating effect sizes and confidence or credible intervals, and use statistics such as *p*-values or Bayes factors as complementary information, rather than as strict thresholds for hypothesis testing (Halsey, 2019).

### Suggestions

Permutation tests can yield correct p-values under appropriate constraints, but, as with parametric models they do not automatically account for non-independent data, and unlike parametric models, they do not account for confounds when estimating effect sizes. For this reason we argue that parametric models could offer a number of important benefits over permutation methods for nodal and dyadic regression. Specifically, well-specified parametric regression models such as OLS or mixed models could be used in the place of node-label permutations. In dyadic regression, multimembership models could be used as an alternative to QAP and its derivatives. In both cases, confounds can be explicitly accounted for without the need to use constrained permutations. These types of models are widely used and have several existing R implementations, for example in MCMCglmm and brms (Hadfield, 2010; Bürkner, 2018). Adopting this approach would yield both correct *p*-values and correct effect size estimates, leading to more reliable statistical inferences.

## Conclusion

In this paper we have highlighted that permutation tests are not a panacea for non-independence in network data. We have illustrated that node-label permutations are equivalent to parametric regression for nodal regression analysis, and that multimembership models can control for non-independence in the same way as QAP in dyadic regression analysis. Given their more widespread use across various life science disciplines, we promote the use of standard parametric models for animal social network analysis in the place of node-label permutations and QAP. Further work is required to understand potential sources of non-independence in animal social networks. We believe that the arguments presented in this study open up opportunities to adopt more powerful, versatile, and robust statistical methods in animal social network analysis.

## Supporting information

Simulation code

## Author contributions

JDAH conceived the idea of the manuscript. The arguments were developed and refined by JDAH, MNW, LJNB, and DWF. The simulations were developed by JDAH with input from MNW, LJNB, and DWF. The manuscript the written by JDAH with input from MNW, LJNB, and DWF.

## Acknowledgements

We would like to thank CRAB and CRAB Social Network Club for feedback and discussions on early versions of this work. JDAH acknowledges funding from the Engineering and Physical Sciences Research Council [grant number EP/R513210/1]. LJNB acknowledges funding from a European Research Council Consolidator Grant (FriendOrigins - 864461) and from the National Institutes of Health (R01AG060931, R01MH118203). DWF and MNW acknowledge funding from the Natural Environment Research Council [grant number NE/S010327/1]. DWF also acknowledges funding from the Natural Environment Research Council [grant number NE/S009914/1]. The authors declare no conflict of interest.

## Supplementary Material: Examples for Multimembership Models in brms and MCMCglmm

This notebook is a short example to demonstrate how multimembership models can be used in place of QAP in dyadic regression. This can be done in a few different R packages, here we show how it can be done in both brms and MCMCglmm.

### Loading packages

Load the packages. asnipe is included to demonstrate the methods against MRQAP using the double semi-partialling method (Dekker et al., 2007).

~~~
library(asnipe)
library(MCMCglmm)
library(brms)
~~~

### Simulate dataset

Next we simulate a dataset with 20 nodes where edge weight (association strength) depends on the sexes of pairs of nodes and the age difference between the nodes. Some individuals are also just more likely to form links, and this creates a node dependence. In this imagined scenario, edge weights are more likely between individuals of different sexes with a large age difference.

~~~
set.seed(1)

# Simulate dyadic data with nodal dependence.
num_nodes <- 20
node_dependence <- rnorm(num_nodes)
sexes <- sample(c(1, 2), num_nodes, replace=TRUE)
mean_ages <- c(20, 10)
ages <- rpois(num_nodes, mean_ages[sexes])

sex_diff <- abs(matrix(rep(sexes, num_nodes), num_nodes, num_nodes)
   - t(matrix(rep(sexes, num_nodes), num_nodes, num_nodes)))
age_diff <- abs(matrix(rep(ages, num_nodes), num_nodes, num_nodes)
   - t(matrix(rep(ages, num_nodes), num_nodes, num_nodes)))
dependencies <- matrix(rep(node_dependence, num_nodes), num_nodes, num_nodes)
dependencies <- dependencies + t(dependencies)

error <- matrix(rnorm(num_nodes^2), num_nodes, num_nodes)
error <- error + t(error)

edge <- 0.2 * sex_diff + 0.2 * age_diff + dependencies + error

Y <- edge
X_age <- age_diff
X_sex <- sex_diff
~~~

We now have three matrices: Y holds edge weights, X_age holds age differences between pairs, and X_sex holds sex differences (binary) between pairs. This is the format required to conduct standard QAP/MRQAP.

### MRQAP

Use asnipe to demonstrate a dyadic regression analysis using conventional MRQAP:

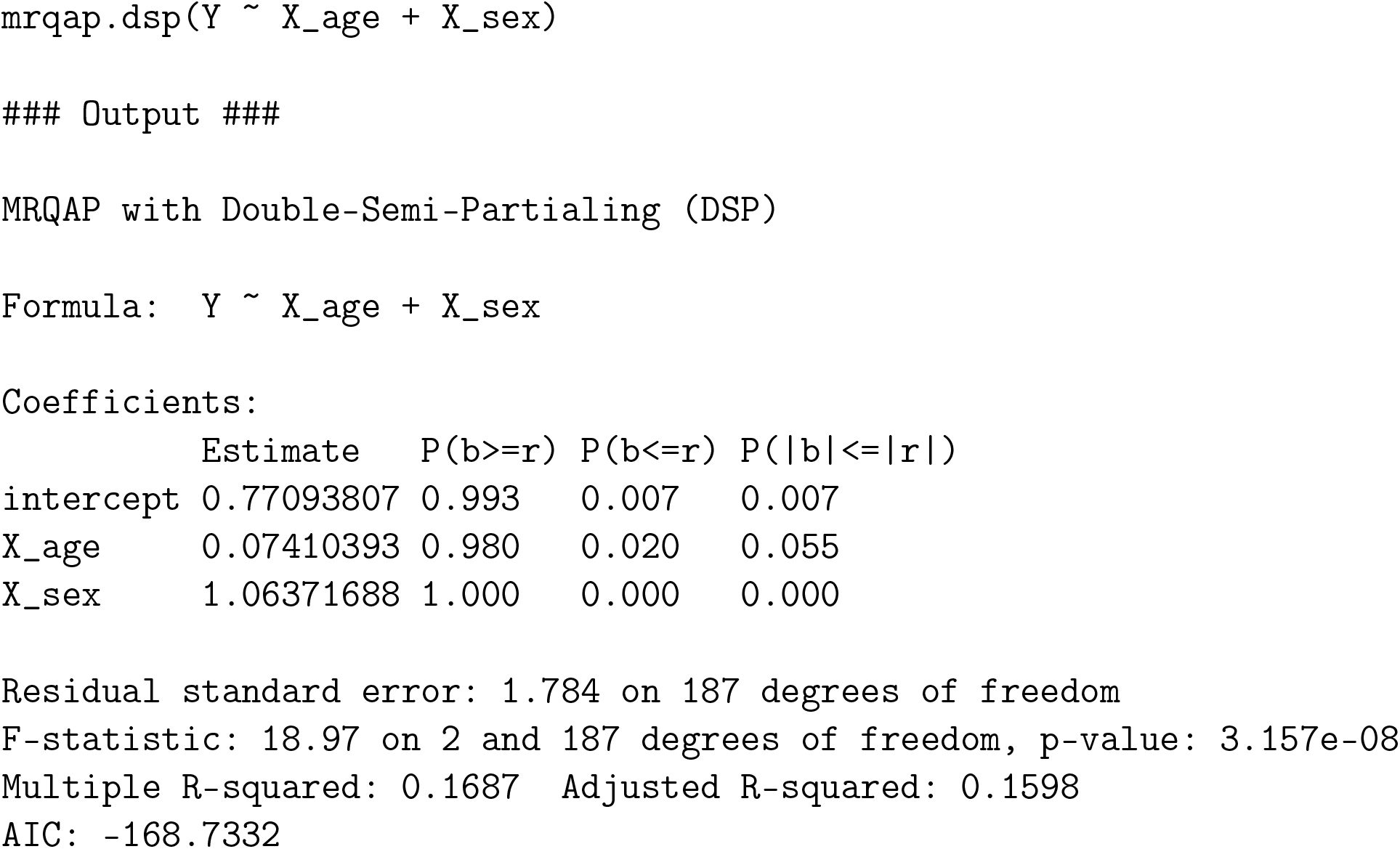

This tells us that age difference is nearly significant and that sex difference is significant.

### Prepare dataframe for brms and MCMCglmm

Now to apply the multimembership model using brms and MCMCglmm. Firstly, both of these packages expect a dataframe, so we need to encourage the data into the correct format. Because our network is undirected, this can be done by taking the upper triangle of each dataframe. We also need to generate a list of node IDs that correspond to the nodes of the network. This is how we will capture the multimembership aspect of the model.

**Note** If working with directed networks, the lower triangle will also need to be included and additional random effects may need to be included to account for influence of a node being a sender or a receiver. These decisions will depend on the data and question, and need to be carefully considered.

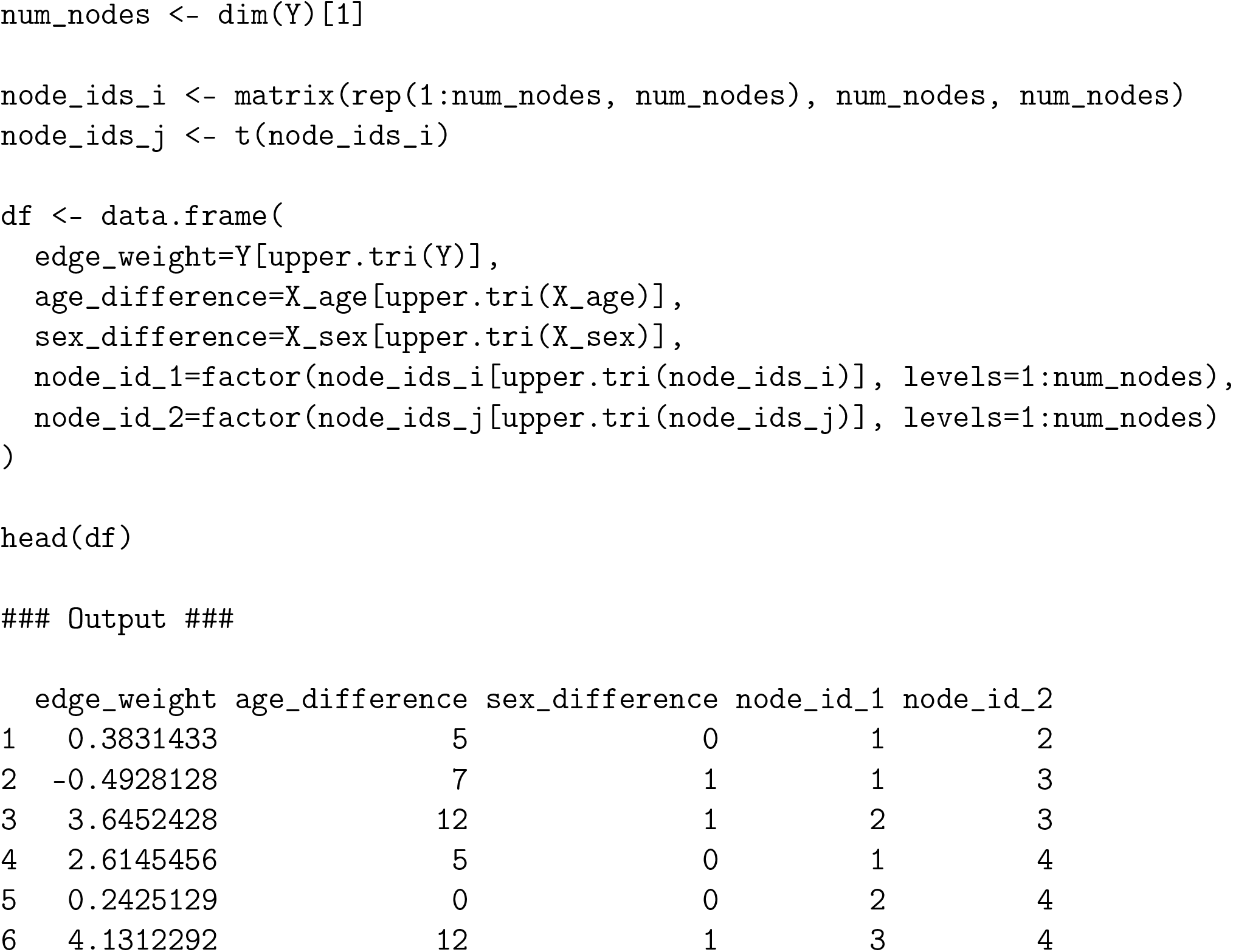

Now we can look at applying multimembership models using MCMCglmm and brms.

### Multimembership models in MCMCglmm

Let’s start with MCMCglmm. The fixed effects part of the model is standard. To include the multimembership part, we use random effects and the mm function with the following notation:

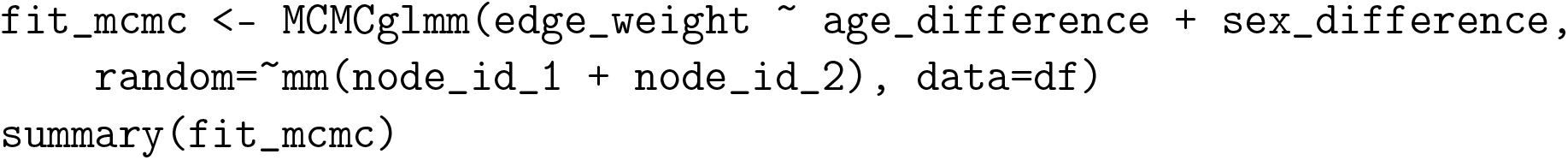
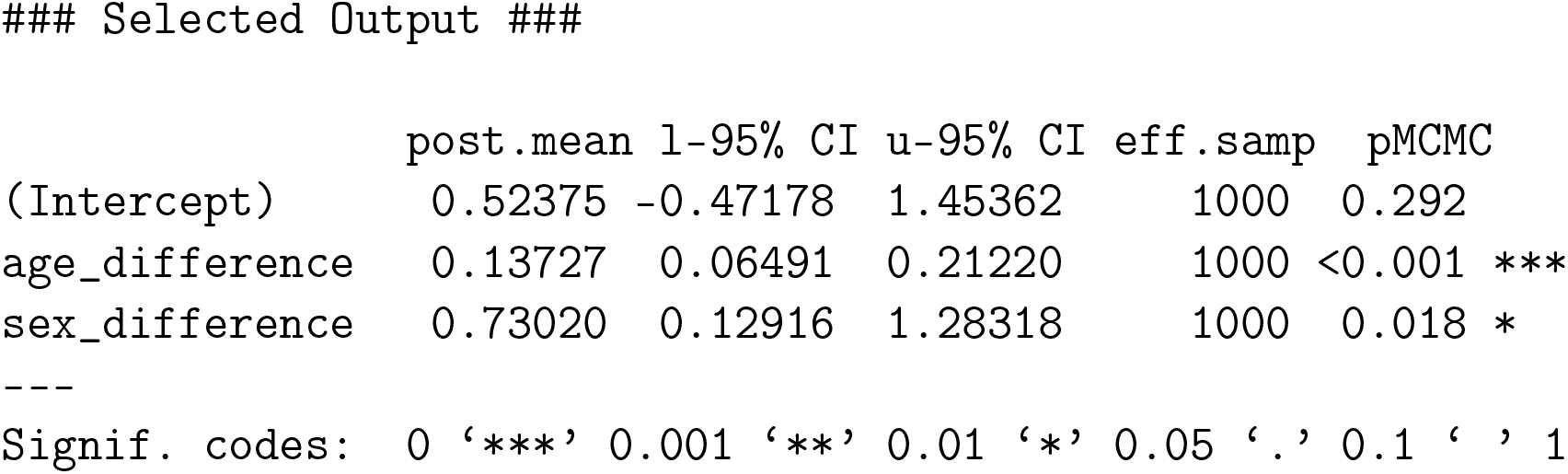

### Multimembership models in brms

brms uses a more conventional notation similar to lme4. Again, the function mm is used to include the multimembership random effects. Note that in this function, the effects are included as separate arguments, separated by a comma, instead of a sum notation like in MCMCglmm.

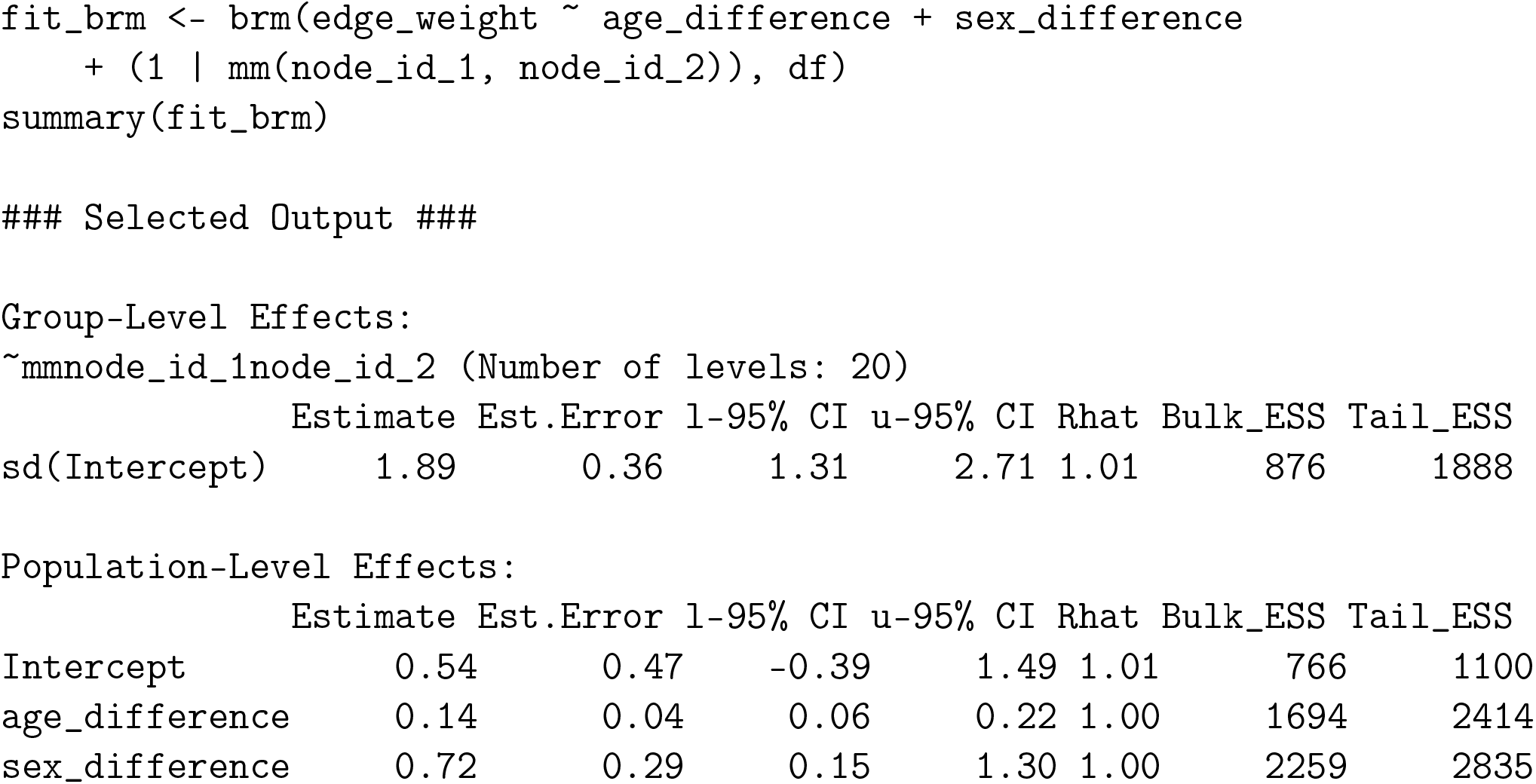

In both of these multimembership models, we see that the coefficient estimates are quite different to those from QAP and have interpretable confidence/credible intervals. This is because these models account for confounds when calculating both the significance (where applicable) and in the effect size estimates, whereas QAP only accounts for confounds when calculating the significance.

## References

Marti J. Anderson and John Robinson. Permutation Tests for Linear Models. Australian & New Zealand Journal of Statistics, 43(1):75–88, 2001. ISSN 1467-842X. doi: 10.1111/1467-842X.00156.

Lars Bejder, David Fletcher, and Stefan Bräger. A method for testing association patterns of social animals. Animal Behaviour, 56(3):719–725, September 1998. ISSN 0003-3472. doi: 10.1006/anbe.1998.0802.

Natasha K. Boyland, David T. Mlynski, Richard James, Lauren J. N. Brent, and Darren P. Croft. The social network structure of a dynamic group of dairy cows: From individual to group level patterns. Applied Animal Behaviour Science, 174:1–10, January 2016. ISSN 0168-1591. doi: 10.1016/j.applanim.2015.11.016.

Carter T. Butts. Social network analysis: A methodological introduction. Asian Journal of Social Psychology, 11(1):13–41, 2008. ISSN 1467-839X. doi: 10.1111/j.1467-839X.2007.00241.x.

Paul-Christian Bürkner. Advanced Bayesian multilevel modeling with the R package brms. The R Journal, 10(1):395–411, 2018. doi: 10.32614/RJ-2018-017.

Jacob Cohen. Statistical Power Analysis. Current Directions in Psychological Science, 1(3):98– 101, June 1992. ISSN 0963-7214. doi: 10.1111/1467-8721.ep10768783.

Jacob Cohen. The earth is round (p<.05). American Psychologist, 49(12):997–1003, 1994. ISSN 1935-990X(Electronic),0003-066X(Print). doi: 10.1037/0003-066X.49.12.997.

Peter Congdon. Bayesian hierarchical models: with applications using R. CRC Press, Taylor & Francis Group, Boca Raton, FL, second edition edition, 2020. ISBN 978-0-429-11335-2.

William L. Cook and David A. Kenny. The Actor–Partner Interdependence Model: A model of bidirectional effects in developmental studies. International Journal of Behavioral Development, 29(2):101–109, March 2005. ISSN 0165-0254. doi: 10.1080/01650250444000405.

Darren P. Croft, Richard James, and Jens Krause. Exploring animal social networks. Princeton University Press, Princeton, N.J., 2010. ISBN 978-1-4008-3776-2.

Darren P. Croft, Joah R. Madden, Daniel W. Franks, and Richard James. Hypothesis testing in animal social networks. Trends in Ecology & Evolution, 26(10):502–507, October 2011. ISSN 01695347. doi: 10.1016/j.tree.2011.05.012.

David Dekker, David Krackhardt, and Tom A. B. Snijders. Sensitivity of MRQAP Tests to Collinearity and Autocorrelation Conditions. Psychometrika, 72(4):563–581, December 2007. ISSN 0033-3123. doi: 10.1007/s11336-007-9016-1.

Dino Dittrich, Roger Th. A. J. Leenders, and Joris Mulder. Network Autocorrelation Modeling: Bayesian Techniques for Estimating and Testing Multiple Network Autocorrelations. Sociological Methodology, 50(1):168–214, August 2020. ISSN 0081-1750. doi: 10.1177/0081175020913899.

Patrick Doreian. Estimating Linear Models with Spatially Distributed Data. Sociological Methodology, 12:359–388, 1981. ISSN 0081-1750. doi: 10.2307/270747.

Norman R. Draper and Harry Smith. Applied Regression Analysis. Wiley Series in Probability and Statistics. Wiley, 1 edition, April 1998. ISBN 978-0-471-17082-2978-1-118-62559-0. doi: 10.1002/9781118625590.

Damien R. Farine. A guide to null models for animal social network analysis. Methods in Ecology and Evolution, 8(10):1309–1320, 2017. ISSN 2041-210X. doi: 10.1111/2041-210X.12772.

Damien R. Farine and Hal Whitehead. Constructing, conducting and interpreting animal social network analysis. Journal of Animal Ecology, 84(5):1144–1163, 2015. ISSN 1365-2656. doi: 10.1111/1365-2656.12418.

Daniel W. Franks, Michael N. Weiss, Matthew J. Silk, Robert J. Y. Perryman, and Darren P. Croft. Calculating effect sizes in animal social network analysis. Methods in Ecology and Evolution, 12(1):33–41, 2021. ISSN 2041-210X. doi: 10.1111/2041-210X.13429.

Phillip I. Good. Permutation tests: a practical guide to resampling methods for testing hypotheses. Springer, New York, 2000. ISBN 978-1-4757-3235-1.

Jarrod D. Hadfield. MCMC methods for multi-response generalized linear mixed models: The MCMCglmm R package. Journal of Statistical Software, 33(2):1–22, 2010.

Lewis G. Halsey. The reign of the p-value is over: what alternative analyses could we employ to fill the power vacuum? Biology Letters, 15(5):20190174, May 2019. doi: 10.1098/rsbl.2019.0174.

David R. Hunter. Curved Exponential Family Models for Social Networks. Social networks, 29 (2):216–230, March 2007. ISSN 0378-8733. doi: 10.1016/j.socnet.2006.08.005.

Sylvia A. Morelli, Desmond C. Ong, Rucha Makati, Matthew O. Jackson, and Jamil Zaki. Empathy and well-being correlate with centrality in different social networks. Proceedings of the National Academy of Sciences, 114(37):9843–9847, September 2017. ISSN 0027-8424, 1091-6490. doi: 10.1073/pnas.1702155114.

Carlo Morselli, Victor Hugo Masias, Fernando Crespo, and Sigifredo Laengle. Predicting sentencing outcomes with centrality measures. Security Informatics, 2(1):4, December 2013. ISSN 2190-8532. doi: 10.1186/2190-8532-2-4.

A. James O’Malley and Peter V. Marsden. The Analysis of Social Networks. Health services & outcomes research methodology, 8(4):222–269, December 2008. ISSN 1387-3741. doi: 10.1007/s10742-008-0041-z.

Judea Pearl. An Introduction to Causal Inference. The International Journal of Biostatistics, 6 (2), January 2010. ISSN 1557-4679. doi: 10.2202/1557-4679.1203.

R Core Team. R: A language and environment for statistical computing. 2020.

Julie Rushmore, Damien Caillaud, Leopold Matamba, Rebecca M. Stumpf, Stephen P. Borgatti, and Sonia Altizer. Social network analysis of wild chimpanzees provides insights for predicting infectious disease risk. Journal of Animal Ecology, 82(5):976–986, 2013. ISSN 1365-2656. doi: 10.1111/1365-2656.12088.

Tom A. B. Snijders, Philippa E. Pattison, Garry L. Robins, and Mark S. Handcock. New Specifications for Exponential Random Graph Models. Sociological Methodology, 36(1):99–153, August 2006. ISSN 0081-1750, 1467-9531. doi: 10.1111/j.1467-9531.2006.00176.x.

Sebastian Sosa, Cédric Sueur, and Ivan Puga-Gonzalez. Network measures in animal social network analysis: Their strengths, limits, interpretations and uses. Methods in Ecology and Evolution, 12(1):10–21, January 2021. ISSN 2041-210X, 2041-210X. doi: 10.1111/2041-210X.13366.

Mark Tranmer, David Steel, and William J. Browne. Multiple-membership multiple-classification models for social network and group dependences. Journal of the Royal Statistical Society: Series A (Statistics in Society), 177(2):439–455, 2014. doi: 10.1111/rssa.12021.

Larry Wasserman. All of Statistics: A Concise Course in Statistical Inference. Springer Texts in Statistics. Springer New York, New York, NY, 2004. ISBN 978-1-4419-2322-6 978-0-38721736-9. doi: 10.1007/978-0-387-21736-9.

William W. S. Wei. Time Series Analysis, March 2013. ISBN: 9780199934898.

Anderson M. Winkler, Matthew A. Webster, Diego Vidaurre, Thomas E. Nichols, and Stephen M. Smith. Multi-level block permutation. NeuroImage, 123:253–268, December 2015. ISSN 1053-8119. doi: 10.1016/j.neuroimage.2015.05.092.

