## Supplementary material for "Common Permutation Methods in Animal Social Network Analysis Do Not Control for Non-independence": Simulation code: dyadic_regression.nb.html

Dyadic regression - Dependence on nodes


Code 

- Show All Code
- Hide All Code
- Download Rmd

### Dyadic regression - Dependence on nodes


```
library(ggplot2)

QAP <- function(Y, X) {
  n <- dim(Y)[1]
  Y_ <- Y[upper.tri(Y)]
  X_ <- X[upper.tri(X)]
  obs <- .lm.fit(cbind(rep(1, length(X_)), X_), Y_)$coefficients[2]
  
  null_dist <- sapply(1:1000, function(i) {
    shuffle_rows <- sample(1:n)
    X_ <- as.vector(X[shuffle_rows, shuffle_rows][upper.tri(X)])
    .lm.fit(cbind(rep(1, length(X_)), X_), Y_)$coefficients[2]
  })
  
  list(estimate=obs, p_value=mean(abs(null_dist) > abs(obs)))
}

dyadic_regression <- function(Y, X) {
  ls_dyadreg <- function(par, X, Y) {
    beta <- par[1:2]
    r <- par[3:(3 + n - 1)]
    
    n <- dim(X)[1]
    Y_ <- Y[upper.tri(Y)]
    X_ <- X[upper.tri(X)]
    
    R <- matrix(rep(r, n), n)
    R <- R + t(R)
    R_ <- R[upper.tri(R)]
    
    Y_pred <- beta[1] + beta[2] * X_ + R_
    sum((Y_ - Y_pred)^2)
  }
  n <- dim(X)[1]
  r <- runif(n, min=-1, max=1)
  beta <- c(0, 0)
  target <- function(par) ls_dyadreg(par, X, Y)
  optim_obj <- optim(c(beta, r), target, method="BFGS", hessian=TRUE)
  samples <- MASS::mvrnorm(1e5, optim_obj$par[1:3], solve(optim_obj$hessian[1:3, 1:3]))
  summary_table <- t(apply(samples, 2, function(x) quantile(x, probs=c(0.025, 0.5, 0.975))))
  rownames(summary_table) <- c("Intercept", "Slope", "Sigma")
  summary_table <- signif(summary_table, 2)
  # summary_table
  summary_table <- cbind(summary_table, sapply(1:3, function(i) 2 * min(mean(samples[, i] < 0), mean(samples[, i] > 0))))
  colnames(summary_table)[4] <- "P-value"
  summary_table
}
```


```
mmlm <- function(Y, X) {
  num_nodes <- dim(Y)[1]

  node_ids_i <- matrix(rep(1:num_nodes, num_nodes), num_nodes, num_nodes)
  node_ids_j <- t(node_ids_i)
  
  df <- data.frame(
    y=Y[upper.tri(Y)],
    x=X[upper.tri(X)],
    node_id_1=factor(node_ids_i[upper.tri(node_ids_i)], levels=1:num_nodes),
    node_id_2=factor(node_ids_j[upper.tri(node_ids_j)], levels=1:num_nodes)
  )
  
  fit_mcmc <- MCMCglmm(y ~ x, random=~mm(node_id_1 + node_id_2), data=df, verbose=FALSE)
  summary(fit_mcmc)$solutions
}
```


```
n <- 20
b <- 0.2

results <- data.frame(effect_size=numeric(), p_value=numeric(), method=numeric(), effect=numeric())

for (effect in c(TRUE, FALSE)) {
  for (iter in 1:1000) {
    X_ <- matrix(runif(n^2), n, n)
    X_ <- X_ * upper.tri(X_)
    X_ <- X_ + t(X_)
    
    Y_ <- matrix(runif(n^2), n, n)
    Y_ <- Y_ * upper.tri(Y_)
    Y_ <- Y_ + t(Y_)

    R <- matrix(rep(runif(n), n), n, n)
    S <- matrix(rep(runif(n), n), n, n)
    
    X = R + t(R) + X_
    Y = S + t(S) + Y_
    
    if (effect) {
      Y <- b * X + (1 - b) * Y
    }
    
    x <- X[upper.tri(X)]
    y <- Y[upper.tri(Y)]
    
    obj_lm <- summary(lm(y ~ x))
    obj_perm <- QAP(Y, X)
    obj_dyadreg <- dyadic_regression(Y, X)
    
    effect_lm <- obj_lm$coefficients[2, 1]
    effect_perm <- obj_perm$estimate
    effect_dyadreg <- obj_dyadreg[2, 2]
    
    pval_lm <- obj_lm$coefficients[2, 4]
    pval_perm <- obj_perm$p_value
    pval_dyadreg <- obj_dyadreg[2, 4]
    
    results[nrow(results) + 1, ] <- list(effect_lm, pval_lm, "OLS", as.character(effect))
    results[nrow(results) + 1, ] <- list(effect_perm, pval_perm, "QAP", as.character(effect))
    results[nrow(results) + 1, ] <- list(effect_dyadreg, pval_dyadreg, "OLS + Control", as.character(effect))
  }
}

write.csv(results, "results/results.dyadic.csv")
results
```


```
agg <- aggregate(effect_size ~ method + effect, results.dyadic, function(x) quantile(x, probs = c(0.025, 0.50, 0.975)))
do.call(data.frame, agg)
```


```
aggregate(p_value ~ method + effect, results.dyadic, function(x) mean(x < 0.05))
```


```
agg <- aggregate(effect_size ~ method + effect, results.dyadic, function(x) signif(quantile(x, probs = c(0.025, 0.50, 0.975)), 3))
do.call(data.frame, agg)
```


```
# Proportion of effect sizes with wrong sign.
aggregate(cbind(wrong_sign=effect_size) ~ method + effect, results.dyadic, function(x) mean(x < 0))
```


```
# Proportion of significant results with wrong effect size sign.
aggregate(cbind(wrong_sign=effect_size) ~ method + effect, subset(results.dyadic, p_value < 0.05 & effect == TRUE), function(x) mean(x < 0))
```
