## Supplementary material for "Common Permutation Methods in Animal Social Network Analysis Do Not Control for Non-independence": Simulation code: dyadic_regression_dependence.nb.html

R Notebook


Code 

- Show All Code
- Hide All Code
- Download Rmd

### R Notebook


```
library(igraph)
```


```
Attaching package: ‘igraph’

The following objects are masked from ‘package:MCMCglmm’:

    path, sir

The following objects are masked from ‘package:ape’:

    edges, mst, ring

The following objects are masked from ‘package:stats’:

    decompose, spectrum

The following object is masked from ‘package:base’:

    union
```


```
n <- 20
b <- 0.2

results <- data.frame(effect_size=numeric(), p_value=numeric(), method=numeric(), effect=numeric())

for (effect in c(TRUE, FALSE)) {
  for (iter in 1:100) {
    X_ <- matrix(runif(n^2), n, n)
    X_ <- X_ * upper.tri(X_)
    X_ <- X_ + t(X_)
    
    Y_ <- matrix(runif(n^2), n, n)
    Y_ <- Y_ * upper.tri(Y_)
    Y_ <- Y_ + t(Y_)

    R <- matrix(rep(runif(n), n), n, n)
    S <- matrix(rep(runif(n), n), n, n)
    
    G <- graph_from_adjacency_matrix(Y_, mode="undirected", weighted=TRUE)

    u <- rep(0, n)
    v <- rep(0, n)
    
    C <- cliques(G, min=4, max=4)
    for (i in 1:length(C)) {
      u[C[[i]]] <- runif(length(C[[i]]))
      v[C[[i]]] <- runif(length(C[[i]]))
    }
    
    U <- matrix(rep(u, n), n, n)
    V <- matrix(rep(v, n), n, n)
    
    X = R + t(R) + U + t(U) + X_
    Y = S + t(S) + V + t(V) + Y_
    
    if (effect) {
      Y <- b * X + (1 - b) * Y
    }
    
    x <- X[upper.tri(X)]
    y <- Y[upper.tri(Y)]
    
    obj_lm <- summary(lm(y ~ x))
    obj_perm <- QAP(Y, X)
    obj_dyadreg <- dyadic_regression(Y, X)
    
    effect_lm <- obj_lm$coefficients[2, 1]
    effect_perm <- obj_perm$estimate
    effect_dyadreg <- obj_dyadreg[2, 2]
    
    pval_lm <- obj_lm$coefficients[2, 4]
    pval_perm <- obj_perm$p_value
    pval_dyadreg <- obj_dyadreg[2, 4]
    
    results[nrow(results) + 1, ] <- list(effect_lm, pval_lm, "OLS", as.character(effect))
    results[nrow(results) + 1, ] <- list(effect_perm, pval_perm, "QAP", as.character(effect))
    results[nrow(results) + 1, ] <- list(effect_dyadreg, pval_dyadreg, "OLS + Control", as.character(effect))
  }
}

# write.csv(results, "results/results.dyadic.clique.csv")
results
```


```
results.dyadic.clique <- read.csv("results/results.dyadic.clique.csv")

ggplot(results.dyadic.clique, aes(x=as.factor(method), y=p_value,  fill=as.factor(effect))) +
  geom_violin(scale="width") +
  scale_fill_manual(values = c(cp[3], cp[8])) +
  geom_abline(slope=0, intercept=0.05, linetype="dashed") +
  labs(x="Method", y="p-value", fill="Effect", title = "Dyadic regression - Dependence on substructure") +
  theme_classic()

ggplot(results.dyadic.clique, aes(x=as.factor(method), y=effect_size,  fill=as.factor(effect))) +
  geom_violin(scale="width") +
  scale_fill_manual(values = c(cp[3], cp[8])) +
  geom_abline(slope=0, intercept=0, linetype="dashed") +
  labs(x="Method", y="Effect Size", fill="Effect", title = "Dyadic regression - Dependence on nodes") +
  theme_classic()

ggsave("figures/dyadic.clique.effect.png", width=4, height=3)
```


```
aggregate(p_value ~ method + effect, results.dyadic.clique, function(x) mean(x < 0.05))
```


```
agg <- aggregate(effect_size ~ method + effect, results.dyadic.clique, function(x) signif(quantile(x, probs = c(0.025, 0.50, 0.975)), 3))
do.call(data.frame, agg)
```


```
# Proportion of effect sizes with wrong sign.
aggregate(cbind(wrong_sign=effect_size) ~ method + effect, results.dyadic.clique, function(x) mean(x < 0))
```


```
# Proportion of significant results with wrong effect size sign.
aggregate(cbind(wrong_sign=effect_size) ~ method + effect, subset(results.dyadic.clique, p_value < 0.05 & effect == TRUE), function(x) mean(x < 0))
```
