## Supplementary material for "Common Permutation Methods in Animal Social Network Analysis Do Not Control for Non-independence": Simulation code: example.nb.html

Multi-membership Models for Dyadic Regression


Code 

- Show All Code
- Hide All Code
- Download Rmd

### Multi-membership Models for Dyadic Regression

This notebook is a short example to demonstrate how multi-membership models can be used in place of QAP in dyadic regression. This can be done in a few different R packages, here we show how it can be done in both `brms` and `MCMCglmm`.

#### Loading packages

Load the packages. `asnipe` is included to demonstrate the methods against MRQAP using the double semi-partialling method described by Dekker et al. (2007).


```
library(asnipe)
library(MCMCglmm)
library(brms)
```


Here we simulate a dataset with 20 nodes where edge weight (association strength) depends on the sexes of pairs of nodes and the age difference between the nodes. Some individuals are also just more likely to form links, and this creates a node dependence. In this imagined scenario, edge weights are more likely between individuals of different sexes with a large age difference.


```
set.seed(1)

# Simulate dyadic data with nodal dependence.
num_nodes <- 20
node_dependence <- rnorm(num_nodes)
sexes <- sample(c(1, 2), num_nodes, replace=TRUE)
mean_ages <- c(20, 10)
ages <- rpois(num_nodes, mean_ages[sexes])

sex_diff <- abs(matrix(rep(sexes, num_nodes), num_nodes, num_nodes) - t(matrix(rep(sexes, num_nodes), num_nodes, num_nodes)))
age_diff <- abs(matrix(rep(ages, num_nodes), num_nodes, num_nodes) - t(matrix(rep(ages, num_nodes), num_nodes, num_nodes)))
dependencies <- matrix(rep(node_dependence, num_nodes), num_nodes, num_nodes)
dependencies <- dependencies + t(dependencies)

error <- matrix(rnorm(num_nodes^2), num_nodes, num_nodes)
error <- error + t(error)

edge <- 0.2 * sex_diff + 0.2 * age_diff + dependencies + error

Y <- edge
X_age <- age_diff
X_sex <- sex_diff
```


We now have three matrices: `Y` holds edge weights, `X_age` holds age differences between pairs, and `X_sex` holds sex differences (binary) between pairs.

This is the format required to conduct standard QAP/MRQAP. We’ll use the package `asnipe` to demonstrate a conventional MRQAP:


```
mrqap.dsp(Y ~ X_age + X_sex)
```


```
MRQAP with Double-Semi-Partialing (DSP)

Formula:  Y ~ X_age + X_sex 

Coefficients:
          Estimate   P(β>=r) P(β<=r) P(|β|<=|r|)
intercept 0.77093807 0.993   0.007   0.007      
X_age     0.07410393 0.980   0.020   0.055      
X_sex     1.06371688 1.000   0.000   0.000      

Residual standard error: 1.784 on 187 degrees of freedom
F-statistic: 18.97 on 2 and 187 degrees of freedom, p-value: 3.157e-08 
Multiple R-squared: 0.1687  Adjusted R-squared: 0.1598 
AIC: -168.7332
```


This tells us that age difference is nearly significant and that sex difference is significant. This sounds reasonable.

Now to apply the multi-membership model using `brms` and `MCMCglmm`. Firstly, both of these packages expect a dataframe, so we need to encourage the data into the correct format. Because our network is undirected, this can be done by taking the upper triangle of each dataframe. We also need to generate a list of node IDs that correspond to the nodes of the network. This is how we will capture the multi-membership aspect of the model.

*Note* If working with directed networks, the lower triangle will also need to be included and additional random effects may need to be included to account for influence of a node being a sender or a receiver. These decisions will depend on the data and question, and need to be carefully considered.


```
num_nodes <- dim(Y)[1]

node_ids_i <- matrix(rep(1:num_nodes, num_nodes), num_nodes, num_nodes)
node_ids_j <- t(node_ids_i)

df <- data.frame(
  edge_weight=Y[upper.tri(Y)],
  age_difference=X_age[upper.tri(X_age)],
  sex_difference=X_sex[upper.tri(X_sex)],
  node_id_1=factor(node_ids_i[upper.tri(node_ids_i)], levels=1:num_nodes),
  node_id_2=factor(node_ids_j[upper.tri(node_ids_j)], levels=1:num_nodes)
)

head(df)
```

#### Multi-membership models in `MCMCglmm`

The fixed effects part of the model is standard. To include the multi-membership part, we use random effects and the `mm` function and the following notation:


```
fit_mcmc <- MCMCglmm(edge_weight ~ age_difference + sex_difference, random=~mm(node_id_1 + node_id_2), data=df)
```


```
                       MCMC iteration = 0

                       MCMC iteration = 1000

                       MCMC iteration = 2000

                       MCMC iteration = 3000

                       MCMC iteration = 4000

                       MCMC iteration = 5000

                       MCMC iteration = 6000

                       MCMC iteration = 7000

                       MCMC iteration = 8000

                       MCMC iteration = 9000

                       MCMC iteration = 10000

                       MCMC iteration = 11000

                       MCMC iteration = 12000

                       MCMC iteration = 13000
```


```
summary(fit_mcmc)
```


```
 Iterations = 3001:12991
 Thinning interval  = 10
 Sample size  = 1000 

 DIC: 671.3743 

 G-structure:  ~mm(node_id_1 + node_id_2)
```


```
 R-structure:  ~units
```


```
 Location effects: edge_weight ~ age_difference + sex_difference 

               post.mean l-95% CI u-95% CI eff.samp pMCMC   
(Intercept)      0.54041 -0.38540  1.48062     1000 0.244   
age_difference   0.13698  0.06452  0.22276     1316 0.002 **
sex_difference   0.72345  0.08560  1.28328     1000 0.024 * 
---
Signif. codes:  0 ‘***’ 0.001 ‘**’ 0.01 ‘*’ 0.05 ‘.’ 0.1 ‘ ’ 1
```


```
fit_mcmc <- MCMCglmm(edge_weight ~ age_difference + sex_difference, random=~mm(node_id_1 + node_id_2), data=df, verbose=FALSE)
summary(fit_mcmc)$solutions
```


```
               post.mean    l-95% CI  u-95% CI eff.samp pMCMC
(Intercept)    0.5357370 -0.37897671 1.4795205     1000 0.236
age_difference 0.1358133  0.06116288 0.2158225     1000 0.001
sex_difference 0.7379256  0.19227004 1.3251552     1000 0.012
```

#### Multi-membership models in `brms`

`brms` uses a more conventional notation similar to `lme4`. Again, the function `mm` is used to include the multi-membership random effects. Note that in this function, the effects are included as separate arguments separated by a comma, instead of a sum notation like in `MCMCglmm`.


```
fit_brm <- brm(edge_weight ~ age_difference + sex_difference + (1 | mm(node_id_1, node_id_2)), df)
summary(fit_brm)
```


In both of these multi-membership models, we see that the coefficient estimates are quite different to those from QAP and have interpretable confidence/credible intervals. This is because these models account for confounds when calculating both the significance (where applicable) and in the effect size estimate, whereas QAP only accounts for confounds when calculating the significance.
