## Supplementary material for "Common Permutation Methods in Animal Social Network Analysis Do Not Control for Non-independence": Simulation code: nodal_regression.nb.html

R Notebook


Code 

- Show All Code
- Hide All Code
- Download Rmd

### R Notebook


```
# Load required libraries
library(igraph)
library(sna)
```


```
Loading required package: statnet.common

Attaching package: ‘statnet.common’

The following object is masked from ‘package:base’:

    order

Loading required package: network
network: Classes for Relational Data
Version 1.16.1 created on 2020-10-06.
copyright (c) 2005, Carter T. Butts, University of California-Irvine
                    Mark S. Handcock, University of California -- Los Angeles
                    David R. Hunter, Penn State University
                    Martina Morris, University of Washington
                    Skye Bender-deMoll, University of Washington
 For citation information, type citation("network").
 Type help("network-package") to get started.


Attaching package: ‘network’

The following objects are masked from ‘package:igraph’:

    %c%, %s%, add.edges, add.vertices, delete.edges, delete.vertices, get.edge.attribute, get.edges, get.vertex.attribute, is.bipartite,
    is.directed, list.edge.attributes, list.vertex.attributes, set.edge.attribute, set.vertex.attribute

sna: Tools for Social Network Analysis
Version 2.6 created on 2020-10-5.
copyright (c) 2005, Carter T. Butts, University of California-Irvine
 For citation information, type citation("sna").
 Type help(package="sna") to get started.


Attaching package: ‘sna’

The following objects are masked from ‘package:igraph’:

    betweenness, bonpow, closeness, components, degree, dyad.census, evcent, hierarchy, is.connected, neighborhood, triad.census
```


```
library(asnipe)
library(lme4)
```


```
Loading required package: Matrix
```


```
library(ggplot2)
as.numeric.factor <- function(x) {as.numeric(levels(x))[x]}
cp <- c("#003f5c", "#2f4b7c", "#665191", "#a05195", "#d45087", "#f95d6a", "#ff7c43", "#ffa600")
```


```
node_permutation <- function(y, x) {
  obs_b <- lm(y ~ x)$coefficients[2]
  null_b <- sapply(1:10000, function(i) .lm.fit(cbind(1, x), sample(y))$coefficients[2])
  mean(abs(obs_b) < abs(null_b))
}
```


```
pvals_lm <- rep(0, 1)
pvals_perm <- rep(0, 1)

results <- data.frame(p_value=numeric(), method=numeric(), effect=numeric())

# Set parameters
areas <- 1
N <- 20
sampling_periods <- 100

for (effect in c(TRUE, FALSE)) {
  for (iter in 1:1000) {
    # Generate nodes
    ids <- data.frame(ID=1:(areas*N),AREA=rep(1:areas,each=N),DEG_DIST=NA)
    
    # Generate a degree distribution and normalise it
    for (i in 1:areas) {
    degree_distribution <- rpois(N,3)+i
    ids$DEG_DIST[ids$AREA==i] <- degree_distribution/max(degree_distribution)
    }
    
    # Generate attributes
    # This creates a correlation between sex and underlying degree probability
    if (effect) {
      ids$SEX <- sapply(ids$DEG_DIST,FUN=function(x) { sample(c("M","F"),1,prob=c(x,1-x))})
    } else {
      ids$SEX <- sample(c("M","F"),N*areas,replace=TRUE)
    }
    # Generate probability for each edge (only within area)
    probs <- outer(ids$DEG_DIST, ids$DEG_DIST, "*")*outer(ids$AREA,ids$AREA,"==")
    
    # Create sampling periods, each sample only contains data from one area
    sps <- array(0,c(areas*sampling_periods,N*areas,N*areas))
    
    for (i in 1:areas) {
    sps[((sampling_periods*(i-1))+1):(sampling_periods*i),
    ((N*(i-1))+1):(N*i),((N*(i-1))+1):(N*i)] <- rgraph(N,m=sampling_periods,tprob=probs[((N*(i-1))+1):(N*i),
    ((N*(i-1))+1):(N*i)],mode="graph")
    }
    
    # Generate network
    network <- get_network(sps,data_format="SP")
    rownames(network) <- ids$SEX
    colnames(network) <- ids$SEX
    
    # Calculate degrees
    ids$DEGREE <- degree(network,gmode="graph")
    
    pval_lm <- summary(lm(DEGREE ~ SEX, data=ids))$coefficients[2, 4]
    pval_perm <- node_permutation(ids$DEGREE, as.numeric(factor(ids$SEX)) - 1)
    
    results[nrow(results) + 1, ] <- list(pval_lm, "OLS", as.character(effect))
    results[nrow(results) + 1, ] <- list(pval_perm, "Node-label Permutation", as.character(effect))
  }
}
```


```
Generating  20  x  20  matrix
Generating  20  x  20  matrix
Generating  20  x  20  matrix
Generating  20  x  20  matrix
Generating  20  x  20  matrix
Generating  20  x  20  matrix
Generating  20  x  20  matrix
Generating  20  x  20  matrix
Generating  20  x  20  matrix
Generating  20  x  20  matrix
Generating  20  x  20  matrix
Generating  20  x  20  matrix
Generating  20  x  20  matrix
Generating  20  x  20  matrix
Generating  20  x  20  matrix
Generating  20  x  20  matrix
Generating  20  x  20  matrix
Generating  20  x  20  matrix
Generating  20  x  20  matrix
Generating  20  x  20  matrix
Generating  20  x  20  matrix
Generating  20  x  20  matrix
Generating  20  x  20  matrix
Generating  20  x  20  matrix
Generating  20  x  20  matrix
Generating  20  x  20  matrix
Generating  20  x  20  matrix
Generating  20  x  20  matrix
Generating  20  x  20  matrix
Generating  20  x  20  matrix
Generating  20  x  20  matrix
Generating  20  x  20  matrix
Generating  20  x  20  matrix
Generating  20  x  20  matrix
Generating  20  x  20  matrix
Generating  20  x  20  matrix
Generating  20  x  20  matrix
Generating  20  x  20  matrix
Generating  20  x  20  matrix
Generating  20  x  20  matrix
Generating  20  x  20  matrix
Generating  20  x  20  matrix
Generating  20  x  20  matrix
Generating  20  x  20  matrix
Generating  20  x  20  matrix
Generating  20  x  20  matrix
Generating  20  x  20  matrix
Generating  20  x  20  matrix
Generating  20  x  20  matrix
Generating  20  x  20  matrix
Generating  20  x  20  matrix
Generating  20  x  20  matrix
Generating  20  x  20  matrix
Generating  20  x  20  matrix
Generating  20  x  20  matrix
Generating  20  x  20  matrix
Generating  20  x  20  matrix
Generating  20  x  20  matrix
Generating  20  x  20  matrix
Generating  20  x  20  matrix
Generating  20  x  20  matrix
Generating  20  x  20  matrix
Generating  20  x  20  matrix
Generating  20  x  20  matrix
Generating  20  x  20  matrix
Generating  20  x  20  matrix
Generating  20  x  20  matrix
Generating  20  x  20  matrix
Generating  20  x  20  matrix
Generating  20  x  20  matrix
Generating  20  x  20  matrix
Generating  20  x  20  matrix
Generating  20  x  20  matrix
Generating  20  x  20  matrix
Generating  20  x  20  matrix
Generating  20  x  20  matrix
Generating  20  x  20  matrix
Generating  20  x  20  matrix
Generating  20  x  20  matrix
Generating  20  x  20  matrix
Generating  20  x  20  matrix
Generating  20  x  20  matrix
Generating  20  x  20  matrix
Generating  20  x  20  matrix
Generating  20  x  20  matrix
Generating  20  x  20  matrix
Generating  20  x  20  matrix
Generating  20  x  20  matrix
Generating  20  x  20  matrix
Generating  20  x  20  matrix
Generating  20  x  20  matrix
Generating  20  x  20  matrix
Generating  20  x  20  matrix
Generating  20  x  20  matrix
Generating  20  x  20  matrix
Generating  20  x  20  matrix
Generating  20  x  20  matrix
Generating  20  x  20  matrix
Generating  20  x  20  matrix
Generating  20  x  20  matrix
Generating  20  x  20  matrix
Generating  20  x  20  matrix
Generating  20  x  20  matrix
Generating  20  x  20  matrix
Generating  20  x  20  matrix
Generating  20  x  20  matrix
Generating  20  x  20  matrix
Generating  20  x  20  matrix
Generating  20  x  20  matrix
Generating  20  x  20  matrix
Generating  20  x  20  matrix
Generating  20  x  20  matrix
Generating  20  x  20  matrix
Generating  20  x  20  matrix
Generating  20  x  20  matrix
Generating  20  x  20  matrix
Generating  20  x  20  matrix
Generating  20  x  20  matrix
Generating  20  x  20  matrix
Generating  20  x  20  matrix
Generating  20  x  20  matrix
Generating  20  x  20  matrix
Generating  20  x  20  matrix
Generating  20  x  20  matrix
Generating  20  x  20  matrix
Generating  20  x  20  matrix
Generating  20  x  20  matrix
Generating  20  x  20  matrix
Generating  20  x  20  matrix
Generating  20  x  20  matrix
Generating  20  x  20  matrix
Generating  20  x  20  matrix
Generating  20  x  20  matrix
Generating  20  x  20  matrix
Generating  20  x  20  matrix
Generating  20  x  20  matrix
Generating  20  x  20  matrix
Generating  20  x  20  matrix
Generating  20  x  20  matrix
Generating  20  x  20  matrix
Generating  20  x  20  matrix
Generating  20  x  20  matrix
Generating  20  x  20  matrix
Generating  20  x  20  matrix
Generating  20  x  20  matrix
Generating  20  x  20  matrix
Generating  20  x  20  matrix
Generating  20  x  20  matrix
Generating  20  x  20  matrix
Generating  20  x  20  matrix
Generating  20  x  20  matrix
Generating  20  x  20  matrix
Generating  20  x  20  matrix
Generating  20  x  20  matrix
Generating  20  x  20  matrix
Generating  20  x  20  matrix
Generating  20  x  20  matrix
Generating  20  x  20  matrix
Generating  20  x  20  matrix
Generating  20  x  20  matrix
Generating  20  x  20  matrix
Generating  20  x  20  matrix
Generating  20  x  20  matrix
Generating  20  x  20  matrix
Generating  20  x  20  matrix
Generating  20  x  20  matrix
Generating  20  x  20  matrix
Generating  20  x  20  matrix
Generating  20  x  20  matrix
Generating  20  x  20  matrix
Generating  20  x  20  matrix
Generating  20  x  20  matrix
Generating  20  x  20  matrix
Generating  20  x  20  matrix
Generating  20  x  20  matrix
Generating  20  x  20  matrix
Generating  20  x  20  matrix
Generating  20  x  20  matrix
Generating  20  x  20  matrix
Generating  20  x  20  matrix
Generating  20  x  20  matrix
Generating  20  x  20  matrix
Generating  20  x  20  matrix
Generating  20  x  20  matrix
Generating  20  x  20  matrix
Generating  20  x  20  matrix
Generating  20  x  20  matrix
Generating  20  x  20  matrix
Generating  20  x  20  matrix
Generating  20  x  20  matrix
Generating  20  x  20  matrix
Generating  20  x  20  matrix
Generating  20  x  20  matrix
Generating  20  x  20  matrix
Generating  20  x  20  matrix
Generating  20  x  20  matrix
Generating  20  x  20  matrix
Generating  20  x  20  matrix
Generating  20  x  20  matrix
Generating  20  x  20  matrix
Generating  20  x  20  matrix
Generating  20  x  20  matrix
Generating  20  x  20  matrix
Generating  20  x  20  matrix
Generating  20  x  20  matrix
Generating  20  x  20  matrix
Generating  20  x  20  matrix
Generating  20  x  20  matrix
Generating  20  x  20  matrix
Generating  20  x  20  matrix
Generating  20  x  20  matrix
Generating  20  x  20  matrix
Generating  20  x  20  matrix
Generating  20  x  20  matrix
Generating  20  x  20  matrix
Generating  20  x  20  matrix
Generating  20  x  20  matrix
Generating  20  x  20  matrix
Generating  20  x  20  matrix
Generating  20  x  20  matrix
Generating  20  x  20  matrix
Generating  20  x  20  matrix
Generating  20  x  20  matrix
Generating  20  x  20  matrix
Generating  20  x  20  matrix
Generating  20  x  20  matrix
Generating  20  x  20  matrix
Generating  20  x  20  matrix
Generating  20  x  20  matrix
Generating  20  x  20  matrix
Generating  20  x  20  matrix
Generating  20  x  20  matrix
Generating  20  x  20  matrix
Generating  20  x  20  matrix
Generating  20  x  20  matrix
Generating  20  x  20  matrix
Generating  20  x  20  matrix
Generating  20  x  20  matrix
Generating  20  x  20  matrix
Generating  20  x  20  matrix
Generating  20  x  20  matrix
Generating  20  x  20  matrix
Generating  20  x  20  matrix
Generating  20  x  20  matrix
Generating  20  x  20  matrix
Generating  20  x  20  matrix
Generating  20  x  20  matrix
Generating  20  x  20  matrix
Generating  20  x  20  matrix
Generating  20  x  20  matrix
Generating  20  x  20  matrix
Generating  20  x  20  matrix
Generating  20  x  20  matrix
Generating  20  x  20  matrix
Generating  20  x  20  matrix
Generating  20  x  20  matrix
Generating  20  x  20  matrix
Generating  20  x  20  matrix
Generating  20  x  20  matrix
Generating  20  x  20  matrix
Generating  20  x  20  matrix
Generating  20  x  20  matrix
Generating  20  x  20  matrix
Generating  20  x  20  matrix
Generating  20  x  20  matrix
Generating  20  x  20  matrix
Generating  20  x  20  matrix
Generating  20  x  20  matrix
Generating  20  x  20  matrix
Generating  20  x  20  matrix
Generating  20  x  20  matrix
Generating  20  x  20  matrix
Generating  20  x  20  matrix
Generating  20  x  20  matrix
Generating  20  x  20  matrix
Generating  20  x  20  matrix
Generating  20  x  20  matrix
Generating  20  x  20  matrix
Generating  20  x  20  matrix
Generating  20  x  20  matrix
Generating  20  x  20  matrix
Generating  20  x  20  matrix
Generating  20  x  20  matrix
Generating  20  x  20  matrix
Generating  20  x  20  matrix
Generating  20  x  20  matrix
Generating  20  x  20  matrix
Generating  20  x  20  matrix
Generating  20  x  20  matrix
Generating  20  x  20  matrix
Generating  20  x  20  matrix
Generating  20  x  20  matrix
Generating  20  x  20  matrix
Generating  20  x  20  matrix
Generating  20  x  20  matrix
Generating  20  x  20  matrix
Generating  20  x  20  matrix
Generating  20  x  20  matrix
Generating  20  x  20  matrix
Generating  20  x  20  matrix
Generating  20  x  20  matrix
Generating  20  x  20  matrix
Generating  20  x  20  matrix
Generating  20  x  20  matrix
Generating  20  x  20  matrix
Generating  20  x  20  matrix
Generating  20  x  20  matrix
Generating  20  x  20  matrix
Generating  20  x  20  matrix
Generating  20  x  20  matrix
Generating  20  x  20  matrix
Generating  20  x  20  matrix
Generating  20  x  20  matrix
Generating  20  x  20  matrix
Generating  20  x  20  matrix
Generating  20  x  20  matrix
Generating  20  x  20  matrix
Generating  20  x  20  matrix
Generating  20  x  20  matrix
Generating  20  x  20  matrix
Generating  20  x  20  matrix
Generating  20  x  20  matrix
Generating  20  x  20  matrix
Generating  20  x  20  matrix
Generating  20  x  20  matrix
Generating  20  x  20  matrix
Generating  20  x  20  matrix
Generating  20  x  20  matrix
Generating  20  x  20  matrix
Generating  20  x  20  matrix
Generating  20  x  20  matrix
Generating  20  x  20  matrix
Generating  20  x  20  matrix
Generating  20  x  20  matrix
Generating  20  x  20  matrix
Generating  20  x  20  matrix
Generating  20  x  20  matrix
Generating  20  x  20  matrix
Generating  20  x  20  matrix
Generating  20  x  20  matrix
Generating  20  x  20  matrix
Generating  20  x  20  matrix
Generating  20  x  20  matrix
Generating  20  x  20  matrix
Generating  20  x  20  matrix
Generating  20  x  20  matrix
Generating  20  x  20  matrix
Generating  20  x  20  matrix
Generating  20  x  20  matrix
Generating  20  x  20  matrix
Generating  20  x  20  matrix
Generating  20  x  20  matrix
Generating  20  x  20  matrix
Generating  20  x  20  matrix
Generating  20  x  20  matrix
Generating  20  x  20  matrix
Generating  20  x  20  matrix
Generating  20  x  20  matrix
Generating  20  x  20  matrix
Generating  20  x  20  matrix
Generating  20  x  20  matrix
Generating  20  x  20  matrix
Generating  20  x  20  matrix
Generating  20  x  20  matrix
Generating  20  x  20  matrix
Generating  20  x  20  matrix
Generating  20  x  20  matrix
Generating  20  x  20  matrix
Generating  20  x  20  matrix
Generating  20  x  20  matrix
Generating  20  x  20  matrix
Generating  20  x  20  matrix
Generating  20  x  20  matrix
Generating  20  x  20  matrix
Generating  20  x  20  matrix
Generating  20  x  20  matrix
Generating  20  x  20  matrix
Generating  20  x  20  matrix
Generating  20  x  20  matrix
Generating  20  x  20  matrix
Generating  20  x  20  matrix
Generating  20  x  20  matrix
Generating  20  x  20  matrix
Generating  20  x  20  matrix
Generating  20  x  20  matrix
Generating  20  x  20  matrix
Generating  20  x  20  matrix
Generating  20  x  20  matrix
Generating  20  x  20  matrix
Generating  20  x  20  matrix
Generating  20  x  20  matrix
Generating  20  x  20  matrix
Generating  20  x  20  matrix
Generating  20  x  20  matrix
Generating  20  x  20  matrix
Generating  20  x  20  matrix
Generating  20  x  20  matrix
Generating  20  x  20  matrix
Generating  20  x  20  matrix
Generating  20  x  20  matrix
Generating  20  x  20  matrix
Generating  20  x  20  matrix
Generating  20  x  20  matrix
Generating  20  x  20  matrix
Generating  20  x  20  matrix
Generating  20  x  20  matrix
Generating  20  x  20  matrix
Generating  20  x  20  matrix
Generating  20  x  20  matrix
Generating  20  x  20  matrix
Generating  20  x  20  matrix
Generating  20  x  20  matrix
Generating  20  x  20  matrix
Generating  20  x  20  matrix
Generating  20  x  20  matrix
Generating  20  x  20  matrix
Generating  20  x  20  matrix
Generating  20  x  20  matrix
Generating  20  x  20  matrix
Generating  20  x  20  matrix
Generating  20  x  20  matrix
Generating  20  x  20  matrix
Generating  20  x  20  matrix
Generating  20  x  20  matrix
Generating  20  x  20  matrix
Generating  20  x  20  matrix
Generating  20  x  20  matrix
Generating  20  x  20  matrix
Generating  20  x  20  matrix
Generating  20  x  20  matrix
Generating  20  x  20  matrix
Generating  20  x  20  matrix
Generating  20  x  20  matrix
Generating  20  x  20  matrix
Generating  20  x  20  matrix
Generating  20  x  20  matrix
Generating  20  x  20  matrix
Generating  20  x  20  matrix
Generating  20  x  20  matrix
Generating  20  x  20  matrix
Generating  20  x  20  matrix
Generating  20  x  20  matrix
Generating  20  x  20  matrix
Generating  20  x  20  matrix
Generating  20  x  20  matrix
Generating  20  x  20  matrix
Generating  20  x  20  matrix
Generating  20  x  20  matrix
Generating  20  x  20  matrix
Generating  20  x  20  matrix
Generating  20  x  20  matrix
Generating  20  x  20  matrix
Generating  20  x  20  matrix
Generating  20  x  20  matrix
Generating  20  x  20  matrix
Generating  20  x  20  matrix
Generating  20  x  20  matrix
Generating  20  x  20  matrix
Generating  20  x  20  matrix
Generating  20  x  20  matrix
Generating  20  x  20  matrix
Generating  20  x  20  matrix
Generating  20  x  20  matrix
Generating  20  x  20  matrix
Generating  20  x  20  matrix
Generating  20  x  20  matrix
Generating  20  x  20  matrix
Generating  20  x  20  matrix
Generating  20  x  20  matrix
Generating  20  x  20  matrix
Generating  20  x  20  matrix
Generating  20  x  20  matrix
Generating  20  x  20  matrix
Generating  20  x  20  matrix
Generating  20  x  20  matrix
Generating  20  x  20  matrix
Generating  20  x  20  matrix
Generating  20  x  20  matrix
Generating  20  x  20  matrix
Generating  20  x  20  matrix
Generating  20  x  20  matrix
Generating  20  x  20  matrix
Generating  20  x  20  matrix
Generating  20  x  20  matrix
Generating  20  x  20  matrix
Generating  20  x  20  matrix
Generating  20  x  20  matrix
Generating  20  x  20  matrix
Generating  20  x  20  matrix
Generating  20  x  20  matrix
Generating  20  x  20  matrix
Generating  20  x  20  matrix
Generating  20  x  20  matrix
Generating  20  x  20  matrix
Generating  20  x  20  matrix
Generating  20  x  20  matrix
Generating  20  x  20  matrix
Generating  20  x  20  matrix
Generating  20  x  20  matrix
Generating  20  x  20  matrix
Generating  20  x  20  matrix
Generating  20  x  20  matrix
Generating  20  x  20  matrix
Generating  20  x  20  matrix
Generating  20  x  20  matrix
Generating  20  x  20  matrix
Generating  20  x  20  matrix
Generating  20  x  20  matrix
Generating  20  x  20  matrix
Generating  20  x  20  matrix
Generating  20  x  20  matrix
Generating  20  x  20  matrix
Generating  20  x  20  matrix
Generating  20  x  20  matrix
Generating  20  x  20  matrix
Generating  20  x  20  matrix
Generating  20  x  20  matrix
Generating  20  x  20  matrix
Generating  20  x  20  matrix
Generating  20  x  20  matrix
Generating  20  x  20  matrix
Generating  20  x  20  matrix
Generating  20  x  20  matrix
Generating  20  x  20  matrix
Generating  20  x  20  matrix
Generating  20  x  20  matrix
Generating  20  x  20  matrix
Generating  20  x  20  matrix
Generating  20  x  20  matrix
Generating  20  x  20  matrix
Generating  20  x  20  matrix
Generating  20  x  20  matrix
Generating  20  x  20  matrix
Generating  20  x  20  matrix
Generating  20  x  20  matrix
Generating  20  x  20  matrix
Generating  20  x  20  matrix
Generating  20  x  20  matrix
Generating  20  x  20  matrix
Generating  20  x  20  matrix
Generating  20  x  20  matrix
Generating  20  x  20  matrix
Generating  20  x  20  matrix
Generating  20  x  20  matrix
Generating  20  x  20  matrix
Generating  20  x  20  matrix
Generating  20  x  20  matrix
Generating  20  x  20  matrix
Generating  20  x  20  matrix
Generating  20  x  20  matrix
Generating  20  x  20  matrix
Generating  20  x  20  matrix
Generating  20  x  20  matrix
Generating  20  x  20  matrix
Generating  20  x  20  matrix
Generating  20  x  20  matrix
Generating  20  x  20  matrix
Generating  20  x  20  matrix
Generating  20  x  20  matrix
Generating  20  x  20  matrix
Generating  20  x  20  matrix
Generating  20  x  20  matrix
Generating  20  x  20  matrix
Generating  20  x  20  matrix
Generating  20  x  20  matrix
Generating  20  x  20  matrix
Generating  20  x  20  matrix
Generating  20  x  20  matrix
Generating  20  x  20  matrix
Generating  20  x  20  matrix
Generating  20  x  20  matrix
Generating  20  x  20  matrix
Generating  20  x  20  matrix
Generating  20  x  20  matrix
Generating  20  x  20  matrix
Generating  20  x  20  matrix
Generating  20  x  20  matrix
Generating  20  x  20  matrix
Generating  20  x  20  matrix
Generating  20  x  20  matrix
Generating  20  x  20  matrix
Generating  20  x  20  matrix
Generating  20  x  20  matrix
Generating  20  x  20  matrix
Generating  20  x  20  matrix
Generating  20  x  20  matrix
Generating  20  x  20  matrix
Generating  20  x  20  matrix
Generating  20  x  20  matrix
Generating  20  x  20  matrix
Generating  20  x  20  matrix
Generating  20  x  20  matrix
Generating  20  x  20  matrix
Generating  20  x  20  matrix
Generating  20  x  20  matrix
Generating  20  x  20  matrix
Generating  20  x  20  matrix
Generating  20  x  20  matrix
Generating  20  x  20  matrix
Generating  20  x  20  matrix
Generating  20  x  20  matrix
Generating  20  x  20  matrix
Generating  20  x  20  matrix
Generating  20  x  20  matrix
Generating  20  x  20  matrix
Generating  20  x  20  matrix
Generating  20  x  20  matrix
Generating  20  x  20  matrix
Generating  20  x  20  matrix
Generating  20  x  20  matrix
Generating  20  x  20  matrix
Generating  20  x  20  matrix
Generating  20  x  20  matrix
Generating  20  x  20  matrix
Generating  20  x  20  matrix
Generating  20  x  20  matrix
Generating  20  x  20  matrix
Generating  20  x  20  matrix
Generating  20  x  20  matrix
Generating  20  x  20  matrix
Generating  20  x  20  matrix
Generating  20  x  20  matrix
Generating  20  x  20  matrix
Generating  20  x  20  matrix
Generating  20  x  20  matrix
Generating  20  x  20  matrix
Generating  20  x  20  matrix
Generating  20  x  20  matrix
Generating  20  x  20  matrix
Generating  20  x  20  matrix
Generating  20  x  20  matrix
Generating  20  x  20  matrix
Generating  20  x  20  matrix
Generating  20  x  20  matrix
Generating  20  x  20  matrix
Generating  20  x  20  matrix
Generating  20  x  20  matrix
Generating  20  x  20  matrix
Generating  20  x  20  matrix
Generating  20  x  20  matrix
Generating  20  x  20  matrix
Generating  20  x  20  matrix
Generating  20  x  20  matrix
Generating  20  x  20  matrix
Generating  20  x  20  matrix
Generating  20  x  20  matrix
Generating  20  x  20  matrix
Generating  20  x  20  matrix
Generating  20  x  20  matrix
Generating  20  x  20  matrix
Generating  20  x  20  matrix
Generating  20  x  20  matrix
Generating  20  x  20  matrix
Generating  20  x  20  matrix
Generating  20  x  20  matrix
Generating  20  x  20  matrix
Generating  20  x  20  matrix
Generating  20  x  20  matrix
Generating  20  x  20  matrix
Generating  20  x  20  matrix
Generating  20  x  20  matrix
Generating  20  x  20  matrix
Generating  20  x  20  matrix
Generating  20  x  20  matrix
Generating  20  x  20  matrix
Generating  20  x  20  matrix
Generating  20  x  20  matrix
Generating  20  x  20  matrix
Generating  20  x  20  matrix
Generating  20  x  20  matrix
Generating  20  x  20  matrix
Generating  20  x  20  matrix
Generating  20  x  20  matrix
Generating  20  x  20  matrix
Generating  20  x  20  matrix
Generating  20  x  20  matrix
Generating  20  x  20  matrix
Generating  20  x  20  matrix
Generating  20  x  20  matrix
Generating  20  x  20  matrix
Generating  20  x  20  matrix
Generating  20  x  20  matrix
Generating  20  x  20  matrix
Generating  20  x  20  matrix
Generating  20  x  20  matrix
Generating  20  x  20  matrix
Generating  20  x  20  matrix
Generating  20  x  20  matrix
Generating  20  x  20  matrix
Generating  20  x  20  matrix
Generating  20  x  20  matrix
Generating  20  x  20  matrix
Generating  20  x  20  matrix
Generating  20  x  20  matrix
Generating  20  x  20  matrix
Generating  20  x  20  matrix
Generating  20  x  20  matrix
Generating  20  x  20  matrix
Generating  20  x  20  matrix
Generating  20  x  20  matrix
Generating  20  x  20  matrix
Generating  20  x  20  matrix
Generating  20  x  20  matrix
Generating  20  x  20  matrix
Generating  20  x  20  matrix
Generating  20  x  20  matrix
Generating  20  x  20  matrix
Generating  20  x  20  matrix
Generating  20  x  20  matrix
Generating  20  x  20  matrix
Generating  20  x  20  matrix
Generating  20  x  20  matrix
Generating  20  x  20  matrix
Generating  20  x  20  matrix
Generating  20  x  20  matrix
Generating  20  x  20  matrix
Generating  20  x  20  matrix
Generating  20  x  20  matrix
Generating  20  x  20  matrix
Generating  20  x  20  matrix
Generating  20  x  20  matrix
Generating  20  x  20  matrix
Generating  20  x  20  matrix
Generating  20  x  20  matrix
Generating  20  x  20  matrix
Generating  20  x  20  matrix
Generating  20  x  20  matrix
Generating  20  x  20  matrix
Generating  20  x  20  matrix
Generating  20  x  20  matrix
Generating  20  x  20  matrix
Generating  20  x  20  matrix
Generating  20  x  20  matrix
Generating  20  x  20  matrix
Generating  20  x  20  matrix
Generating  20  x  20  matrix
Generating  20  x  20  matrix
Generating  20  x  20  matrix
Generating  20  x  20  matrix
Generating  20  x  20  matrix
Generating  20  x  20  matrix
Generating  20  x  20  matrix
Generating  20  x  20  matrix
Generating  20  x  20  matrix
Generating  20  x  20  matrix
Generating  20  x  20  matrix
Generating  20  x  20  matrix
Generating  20  x  20  matrix
Generating  20  x  20  matrix
Generating  20  x  20  matrix
Generating  20  x  20  matrix
Generating  20  x  20  matrix
Generating  20  x  20  matrix
Generating  20  x  20  matrix
Generating  20  x  20  matrix
Generating  20  x  20  matrix
Generating  20  x  20  matrix
Generating  20  x  20  matrix
Generating  20  x  20  matrix
Generating  20  x  20  matrix
Generating  20  x  20  matrix
Generating  20  x  20  matrix
Generating  20  x  20  matrix
Generating  20  x  20  matrix
Generating  20  x  20  matrix
Generating  20  x  20  matrix
Generating  20  x  20  matrix
Generating  20  x  20  matrix
Generating  20  x  20  matrix
Generating  20  x  20  matrix
Generating  20  x  20  matrix
Generating  20  x  20  matrix
Generating  20  x  20  matrix
Generating  20  x  20  matrix
Generating  20  x  20  matrix
Generating  20  x  20  matrix
Generating  20  x  20  matrix
Generating  20  x  20  matrix
Generating  20  x  20  matrix
Generating  20  x  20  matrix
Generating  20  x  20  matrix
Generating  20  x  20  matrix
Generating  20  x  20  matrix
Generating  20  x  20  matrix
Generating  20  x  20  matrix
Generating  20  x  20  matrix
Generating  20  x  20  matrix
Generating  20  x  20  matrix
Generating  20  x  20  matrix
Generating  20  x  20  matrix
Generating  20  x  20  matrix
Generating  20  x  20  matrix
Generating  20  x  20  matrix
Generating  20  x  20  matrix
Generating  20  x  20  matrix
Generating  20  x  20  matrix
Generating  20  x  20  matrix
Generating  20  x  20  matrix
Generating  20  x  20  matrix
Generating  20  x  20  matrix
Generating  20  x  20  matrix
Generating  20  x  20  matrix
Generating  20  x  20  matrix
Generating  20  x  20  matrix
Generating  20  x  20  matrix
Generating  20  x  20  matrix
Generating  20  x  20  matrix
Generating  20  x  20  matrix
Generating  20  x  20  matrix
Generating  20  x  20  matrix
Generating  20  x  20  matrix
Generating  20  x  20  matrix
Generating  20  x  20  matrix
Generating  20  x  20  matrix
Generating  20  x  20  matrix
Generating  20  x  20  matrix
Generating  20  x  20  matrix
Generating  20  x  20  matrix
Generating  20  x  20  matrix
Generating  20  x  20  matrix
Generating  20  x  20  matrix
Generating  20  x  20  matrix
Generating  20  x  20  matrix
Generating  20  x  20  matrix
Generating  20  x  20  matrix
Generating  20  x  20  matrix
Generating  20  x  20  matrix
Generating  20  x  20  matrix
Generating  20  x  20  matrix
Generating  20  x  20  matrix
Generating  20  x  20  matrix
Generating  20  x  20  matrix
Generating  20  x  20  matrix
Generating  20  x  20  matrix
Generating  20  x  20  matrix
Generating  20  x  20  matrix
Generating  20  x  20  matrix
Generating  20  x  20  matrix
Generating  20  x  20  matrix
Generating  20  x  20  matrix
Generating  20  x  20  matrix
Generating  20  x  20  matrix
Generating  20  x  20  matrix
Generating  20  x  20  matrix
Generating  20  x  20  matrix
Generating  20  x  20  matrix
Generating  20  x  20  matrix
Generating  20  x  20  matrix
Generating  20  x  20  matrix
Generating  20  x  20  matrix
Generating  20  x  20  matrix
Generating  20  x  20  matrix
Generating  20  x  20  matrix
Generating  20  x  20  matrix
Generating  20  x  20  matrix
Generating  20  x  20  matrix
Generating  20  x  20  matrix
Generating  20  x  20  matrix
Generating  20  x  20  matrix
Generating  20  x  20  matrix
Generating  20  x  20  matrix
Generating  20  x  20  matrix
Generating  20  x  20  matrix
Generating  20  x  20  matrix
Generating  20  x  20  matrix
Generating  20  x  20  matrix
Generating  20  x  20  matrix
Generating  20  x  20  matrix
Generating  20  x  20  matrix
Generating  20  x  20  matrix
Generating  20  x  20  matrix
Generating  20  x  20  matrix
Generating  20  x  20  matrix
Generating  20  x  20  matrix
Generating  20  x  20  matrix
Generating  20  x  20  matrix
Generating  20  x  20  matrix
Generating  20  x  20  matrix
Generating  20  x  20  matrix
Generating  20  x  20  matrix
Generating  20  x  20  matrix
Generating  20  x  20  matrix
Generating  20  x  20  matrix
Generating  20  x  20  matrix
Generating  20  x  20  matrix
Generating  20  x  20  matrix
Generating  20  x  20  matrix
Generating  20  x  20  matrix
Generating  20  x  20  matrix
Generating  20  x  20  matrix
Generating  20  x  20  matrix
Generating  20  x  20  matrix
Generating  20  x  20  matrix
Generating  20  x  20  matrix
Generating  20  x  20  matrix
Generating  20  x  20  matrix
Generating  20  x  20  matrix
Generating  20  x  20  matrix
Generating  20  x  20  matrix
Generating  20  x  20  matrix
Generating  20  x  20  matrix
Generating  20  x  20  matrix
Generating  20  x  20  matrix
Generating  20  x  20  matrix
Generating  20  x  20  matrix
Generating  20  x  20  matrix
Generating  20  x  20  matrix
Generating  20  x  20  matrix
Generating  20  x  20  matrix
Generating  20  x  20  matrix
Generating  20  x  20  matrix
Generating  20  x  20  matrix
Generating  20  x  20  matrix
Generating  20  x  20  matrix
Generating  20  x  20  matrix
Generating  20  x  20  matrix
Generating  20  x  20  matrix
Generating  20  x  20  matrix
Generating  20  x  20  matrix
Generating  20  x  20  matrix
Generating  20  x  20  matrix
Generating  20  x  20  matrix
Generating  20  x  20  matrix
Generating  20  x  20  matrix
Generating  20  x  20  matrix
Generating  20  x  20  matrix
Generating  20  x  20  matrix
Generating  20  x  20  matrix
Generating  20  x  20  matrix
Generating  20  x  20  matrix
Generating  20  x  20  matrix
Generating  20  x  20  matrix
Generating  20  x  20  matrix
Generating  20  x  20  matrix
Generating  20  x  20  matrix
Generating  20  x  20  matrix
Generating  20  x  20  matrix
Generating  20  x  20  matrix
Generating  20  x  20  matrix
Generating  20  x  20  matrix
Generating  20  x  20  matrix
Generating  20  x  20  matrix
Generating  20  x  20  matrix
Generating  20  x  20  matrix
Generating  20  x  20  matrix
Generating  20  x  20  matrix
Generating  20  x  20  matrix
Generating  20  x  20  matrix
Generating  20  x  20  matrix
Generating  20  x  20  matrix
Generating  20  x  20  matrix
Generating  20  x  20  matrix
Generating  20  x  20  matrix
Generating  20  x  20  matrix
Generating  20  x  20  matrix
Generating  20  x  20  matrix
Generating  20  x  20  matrix
Generating  20  x  20  matrix
Generating  20  x  20  matrix
Generating  20  x  20  matrix
Generating  20  x  20  matrix
Generating  20  x  20  matrix
Generating  20  x  20  matrix
Generating  20  x  20  matrix
Generating  20  x  20  matrix
Generating  20  x  20  matrix
Generating  20  x  20  matrix
Generating  20  x  20  matrix
Generating  20  x  20  matrix
Generating  20  x  20  matrix
Generating  20  x  20  matrix
Generating  20  x  20  matrix
Generating  20  x  20  matrix
Generating  20  x  20  matrix
Generating  20  x  20  matrix
Generating  20  x  20  matrix
Generating  20  x  20  matrix
Generating  20  x  20  matrix
Generating  20  x  20  matrix
Generating  20  x  20  matrix
Generating  20  x  20  matrix
Generating  20  x  20  matrix
Generating  20  x  20  matrix
Generating  20  x  20  matrix
Generating  20  x  20  matrix
Generating  20  x  20  matrix
Generating  20  x  20  matrix
Generating  20  x  20  matrix
Generating  20  x  20  matrix
Generating  20  x  20  matrix
Generating  20  x  20  matrix
Generating  20  x  20  matrix
Generating  20  x  20  matrix
Generating  20  x  20  matrix
Generating  20  x  20  matrix
Generating  20  x  20  matrix
Generating  20  x  20  matrix
Generating  20  x  20  matrix
Generating  20  x  20  matrix
Generating  20  x  20  matrix
Generating  20  x  20  matrix
Generating  20  x  20  matrix
Generating  20  x  20  matrix
Generating  20  x  20  matrix
Generating  20  x  20  matrix
Generating  20  x  20  matrix
Generating  20  x  20  matrix
Generating  20  x  20  matrix
Generating  20  x  20  matrix
Generating  20  x  20  matrix
Generating  20  x  20  matrix
Generating  20  x  20  matrix
Generating  20  x  20  matrix
Generating  20  x  20  matrix
Generating  20  x  20  matrix
Generating  20  x  20  matrix
Generating  20  x  20  matrix
Generating  20  x  20  matrix
Generating  20  x  20  matrix
Generating  20  x  20  matrix
Generating  20  x  20  matrix
Generating  20  x  20  matrix
Generating  20  x  20  matrix
Generating  20  x  20  matrix
Generating  20  x  20  matrix
Generating  20  x  20  matrix
Generating  20  x  20  matrix
Generating  20  x  20  matrix
Generating  20  x  20  matrix
Generating  20  x  20  matrix
Generating  20  x  20  matrix
Generating  20  x  20  matrix
Generating  20  x  20  matrix
Generating  20  x  20  matrix
Generating  20  x  20  matrix
Generating  20  x  20  matrix
Generating  20  x  20  matrix
Generating  20  x  20  matrix
Generating  20  x  20  matrix
Generating  20  x  20  matrix
Generating  20  x  20  matrix
Generating  20  x  20  matrix
Generating  20  x  20  matrix
Generating  20  x  20  matrix
Generating  20  x  20  matrix
Generating  20  x  20  matrix
Generating  20  x  20  matrix
Generating  20  x  20  matrix
Generating  20  x  20  matrix
Generating  20  x  20  matrix
Generating  20  x  20  matrix
Generating  20  x  20  matrix
Generating  20  x  20  matrix
Generating  20  x  20  matrix
Generating  20  x  20  matrix
Generating  20  x  20  matrix
Generating  20  x  20  matrix
Generating  20  x  20  matrix
Generating  20  x  20  matrix
Generating  20  x  20  matrix
Generating  20  x  20  matrix
Generating  20  x  20  matrix
Generating  20  x  20  matrix
Generating  20  x  20  matrix
Generating  20  x  20  matrix
Generating  20  x  20  matrix
Generating  20  x  20  matrix
Generating  20  x  20  matrix
Generating  20  x  20  matrix
Generating  20  x  20  matrix
Generating  20  x  20  matrix
Generating  20  x  20  matrix
Generating  20  x  20  matrix
Generating  20  x  20  matrix
Generating  20  x  20  matrix
Generating  20  x  20  matrix
Generating  20  x  20  matrix
Generating  20  x  20  matrix
Generating  20  x  20  matrix
Generating  20  x  20  matrix
Generating  20  x  20  matrix
Generating  20  x  20  matrix
Generating  20  x  20  matrix
Generating  20  x  20  matrix
Generating  20  x  20  matrix
Generating  20  x  20  matrix
Generating  20  x  20  matrix
Generating  20  x  20  matrix
Generating  20  x  20  matrix
Generating  20  x  20  matrix
Generating  20  x  20  matrix
Generating  20  x  20  matrix
Generating  20  x  20  matrix
Generating  20  x  20  matrix
Generating  20  x  20  matrix
Generating  20  x  20  matrix
Generating  20  x  20  matrix
Generating  20  x  20  matrix
Generating  20  x  20  matrix
Generating  20  x  20  matrix
Generating  20  x  20  matrix
Generating  20  x  20  matrix
Generating  20  x  20  matrix
Generating  20  x  20  matrix
Generating  20  x  20  matrix
Generating  20  x  20  matrix
Generating  20  x  20  matrix
Generating  20  x  20  matrix
Generating  20  x  20  matrix
Generating  20  x  20  matrix
Generating  20  x  20  matrix
Generating  20  x  20  matrix
Generating  20  x  20  matrix
Generating  20  x  20  matrix
Generating  20  x  20  matrix
Generating  20  x  20  matrix
Generating  20  x  20  matrix
Generating  20  x  20  matrix
Generating  20  x  20  matrix
Generating  20  x  20  matrix
Generating  20  x  20  matrix
Generating  20  x  20  matrix
Generating  20  x  20  matrix
Generating  20  x  20  matrix
Generating  20  x  20  matrix
Generating  20  x  20  matrix
Generating  20  x  20  matrix
Generating  20  x  20  matrix
Generating  20  x  20  matrix
Generating  20  x  20  matrix
Generating  20  x  20  matrix
Generating  20  x  20  matrix
Generating  20  x  20  matrix
Generating  20  x  20  matrix
Generating  20  x  20  matrix
Generating  20  x  20  matrix
Generating  20  x  20  matrix
Generating  20  x  20  matrix
Generating  20  x  20  matrix
Generating  20  x  20  matrix
Generating  20  x  20  matrix
Generating  20  x  20  matrix
Generating  20  x  20  matrix
Generating  20  x  20  matrix
Generating  20  x  20  matrix
Generating  20  x  20  matrix
Generating  20  x  20  matrix
Generating  20  x  20  matrix
Generating  20  x  20  matrix
Generating  20  x  20  matrix
Generating  20  x  20  matrix
Generating  20  x  20  matrix
Generating  20  x  20  matrix
Generating  20  x  20  matrix
Generating  20  x  20  matrix
Generating  20  x  20  matrix
Generating  20  x  20  matrix
Generating  20  x  20  matrix
Generating  20  x  20  matrix
Generating  20  x  20  matrix
Generating  20  x  20  matrix
Generating  20  x  20  matrix
Generating  20  x  20  matrix
Generating  20  x  20  matrix
Generating  20  x  20  matrix
Generating  20  x  20  matrix
Generating  20  x  20  matrix
Generating  20  x  20  matrix
Generating  20  x  20  matrix
Generating  20  x  20  matrix
Generating  20  x  20  matrix
Generating  20  x  20  matrix
Generating  20  x  20  matrix
Generating  20  x  20  matrix
Generating  20  x  20  matrix
Generating  20  x  20  matrix
Generating  20  x  20  matrix
Generating  20  x  20  matrix
Generating  20  x  20  matrix
Generating  20  x  20  matrix
Generating  20  x  20  matrix
Generating  20  x  20  matrix
Generating  20  x  20  matrix
Generating  20  x  20  matrix
Generating  20  x  20  matrix
Generating  20  x  20  matrix
Generating  20  x  20  matrix
Generating  20  x  20  matrix
Generating  20  x  20  matrix
Generating  20  x  20  matrix
Generating  20  x  20  matrix
Generating  20  x  20  matrix
Generating  20  x  20  matrix
Generating  20  x  20  matrix
Generating  20  x  20  matrix
Generating  20  x  20  matrix
Generating  20  x  20  matrix
Generating  20  x  20  matrix
Generating  20  x  20  matrix
Generating  20  x  20  matrix
Generating  20  x  20  matrix
Generating  20  x  20  matrix
Generating  20  x  20  matrix
Generating  20  x  20  matrix
Generating  20  x  20  matrix
Generating  20  x  20  matrix
Generating  20  x  20  matrix
Generating  20  x  20  matrix
Generating  20  x  20  matrix
Generating  20  x  20  matrix
Generating  20  x  20  matrix
Generating  20  x  20  matrix
Generating  20  x  20  matrix
Generating  20  x  20  matrix
Generating  20  x  20  matrix
Generating  20  x  20  matrix
Generating  20  x  20  matrix
Generating  20  x  20  matrix
Generating  20  x  20  matrix
Generating  20  x  20  matrix
Generating  20  x  20  matrix
Generating  20  x  20  matrix
Generating  20  x  20  matrix
Generating  20  x  20  matrix
Generating  20  x  20  matrix
Generating  20  x  20  matrix
Generating  20  x  20  matrix
Generating  20  x  20  matrix
Generating  20  x  20  matrix
Generating  20  x  20  matrix
Generating  20  x  20  matrix
Generating  20  x  20  matrix
Generating  20  x  20  matrix
Generating  20  x  20  matrix
Generating  20  x  20  matrix
Generating  20  x  20  matrix
Generating  20  x  20  matrix
Generating  20  x  20  matrix
Generating  20  x  20  matrix
Generating  20  x  20  matrix
Generating  20  x  20  matrix
Generating  20  x  20  matrix
Generating  20  x  20  matrix
Generating  20  x  20  matrix
Generating  20  x  20  matrix
Generating  20  x  20  matrix
Generating  20  x  20  matrix
Generating  20  x  20  matrix
Generating  20  x  20  matrix
Generating  20  x  20  matrix
Generating  20  x  20  matrix
Generating  20  x  20  matrix
Generating  20  x  20  matrix
Generating  20  x  20  matrix
Generating  20  x  20  matrix
Generating  20  x  20  matrix
Generating  20  x  20  matrix
Generating  20  x  20  matrix
Generating  20  x  20  matrix
Generating  20  x  20  matrix
Generating  20  x  20  matrix
Generating  20  x  20  matrix
Generating  20  x  20  matrix
Generating  20  x  20  matrix
Generating  20  x  20  matrix
Generating  20  x  20  matrix
Generating  20  x  20  matrix
Generating  20  x  20  matrix
Generating  20  x  20  matrix
Generating  20  x  20  matrix
Generating  20  x  20  matrix
Generating  20  x  20  matrix
Generating  20  x  20  matrix
Generating  20  x  20  matrix
Generating  20  x  20  matrix
Generating  20  x  20  matrix
Generating  20  x  20  matrix
Generating  20  x  20  matrix
Generating  20  x  20  matrix
Generating  20  x  20  matrix
Generating  20  x  20  matrix
Generating  20  x  20  matrix
Generating  20  x  20  matrix
Generating  20  x  20  matrix
Generating  20  x  20  matrix
Generating  20  x  20  matrix
Generating  20  x  20  matrix
Generating  20  x  20  matrix
Generating  20  x  20  matrix
Generating  20  x  20  matrix
Generating  20  x  20  matrix
Generating  20  x  20  matrix
Generating  20  x  20  matrix
Generating  20  x  20  matrix
Generating  20  x  20  matrix
Generating  20  x  20  matrix
Generating  20  x  20  matrix
Generating  20  x  20  matrix
Generating  20  x  20  matrix
Generating  20  x  20  matrix
Generating  20  x  20  matrix
Generating  20  x  20  matrix
Generating  20  x  20  matrix
Generating  20  x  20  matrix
Generating  20  x  20  matrix
Generating  20  x  20  matrix
Generating  20  x  20  matrix
Generating  20  x  20  matrix
Generating  20  x  20  matrix
Generating  20  x  20  matrix
Generating  20  x  20  matrix
Generating  20  x  20  matrix
Generating  20  x  20  matrix
Generating  20  x  20  matrix
Generating  20  x  20  matrix
Generating  20  x  20  matrix
Generating  20  x  20  matrix
Generating  20  x  20  matrix
Generating  20  x  20  matrix
Generating  20  x  20  matrix
Generating  20  x  20  matrix
Generating  20  x  20  matrix
Generating  20  x  20  matrix
Generating  20  x  20  matrix
Generating  20  x  20  matrix
Generating  20  x  20  matrix
Generating  20  x  20  matrix
Generating  20  x  20  matrix
Generating  20  x  20  matrix
Generating  20  x  20  matrix
Generating  20  x  20  matrix
Generating  20  x  20  matrix
Generating  20  x  20  matrix
Generating  20  x  20  matrix
Generating  20  x  20  matrix
Generating  20  x  20  matrix
Generating  20  x  20  matrix
Generating  20  x  20  matrix
Generating  20  x  20  matrix
Generating  20  x  20  matrix
Generating  20  x  20  matrix
Generating  20  x  20  matrix
Generating  20  x  20  matrix
Generating  20  x  20  matrix
Generating  20  x  20  matrix
Generating  20  x  20  matrix
Generating  20  x  20  matrix
Generating  20  x  20  matrix
Generating  20  x  20  matrix
Generating  20  x  20  matrix
Generating  20  x  20  matrix
Generating  20  x  20  matrix
Generating  20  x  20  matrix
Generating  20  x  20  matrix
Generating  20  x  20  matrix
Generating  20  x  20  matrix
Generating  20  x  20  matrix
Generating  20  x  20  matrix
Generating  20  x  20  matrix
Generating  20  x  20  matrix
Generating  20  x  20  matrix
Generating  20  x  20  matrix
Generating  20  x  20  matrix
Generating  20  x  20  matrix
Generating  20  x  20  matrix
Generating  20  x  20  matrix
Generating  20  x  20  matrix
Generating  20  x  20  matrix
Generating  20  x  20  matrix
Generating  20  x  20  matrix
Generating  20  x  20  matrix
Generating  20  x  20  matrix
Generating  20  x  20  matrix
Generating  20  x  20  matrix
Generating  20  x  20  matrix
Generating  20  x  20  matrix
Generating  20  x  20  matrix
Generating  20  x  20  matrix
Generating  20  x  20  matrix
Generating  20  x  20  matrix
Generating  20  x  20  matrix
Generating  20  x  20  matrix
Generating  20  x  20  matrix
Generating  20  x  20  matrix
Generating  20  x  20  matrix
Generating  20  x  20  matrix
Generating  20  x  20  matrix
Generating  20  x  20  matrix
Generating  20  x  20  matrix
Generating  20  x  20  matrix
Generating  20  x  20  matrix
Generating  20  x  20  matrix
Generating  20  x  20  matrix
Generating  20  x  20  matrix
Generating  20  x  20  matrix
Generating  20  x  20  matrix
Generating  20  x  20  matrix
Generating  20  x  20  matrix
Generating  20  x  20  matrix
Generating  20  x  20  matrix
Generating  20  x  20  matrix
Generating  20  x  20  matrix
Generating  20  x  20  matrix
Generating  20  x  20  matrix
Generating  20  x  20  matrix
Generating  20  x  20  matrix
Generating  20  x  20  matrix
Generating  20  x  20  matrix
Generating  20  x  20  matrix
Generating  20  x  20  matrix
Generating  20  x  20  matrix
Generating  20  x  20  matrix
Generating  20  x  20  matrix
Generating  20  x  20  matrix
Generating  20  x  20  matrix
Generating  20  x  20  matrix
Generating  20  x  20  matrix
Generating  20  x  20  matrix
Generating  20  x  20  matrix
Generating  20  x  20  matrix
Generating  20  x  20  matrix
Generating  20  x  20  matrix
Generating  20  x  20  matrix
Generating  20  x  20  matrix
Generating  20  x  20  matrix
Generating  20  x  20  matrix
Generating  20  x  20  matrix
Generating  20  x  20  matrix
Generating  20  x  20  matrix
Generating  20  x  20  matrix
Generating  20  x  20  matrix
Generating  20  x  20  matrix
Generating  20  x  20  matrix
Generating  20  x  20  matrix
Generating  20  x  20  matrix
Generating  20  x  20  matrix
Generating  20  x  20  matrix
Generating  20  x  20  matrix
Generating  20  x  20  matrix
Generating  20  x  20  matrix
Generating  20  x  20  matrix
Generating  20  x  20  matrix
Generating  20  x  20  matrix
Generating  20  x  20  matrix
Generating  20  x  20  matrix
Generating  20  x  20  matrix
Generating  20  x  20  matrix
Generating  20  x  20  matrix
Generating  20  x  20  matrix
Generating  20  x  20  matrix
Generating  20  x  20  matrix
Generating  20  x  20  matrix
Generating  20  x  20  matrix
Generating  20  x  20  matrix
Generating  20  x  20  matrix
Generating  20  x  20  matrix
Generating  20  x  20  matrix
Generating  20  x  20  matrix
Generating  20  x  20  matrix
Generating  20  x  20  matrix
Generating  20  x  20  matrix
Generating  20  x  20  matrix
Generating  20  x  20  matrix
Generating  20  x  20  matrix
Generating  20  x  20  matrix
Generating  20  x  20  matrix
Generating  20  x  20  matrix
Generating  20  x  20  matrix
Generating  20  x  20  matrix
Generating  20  x  20  matrix
Generating  20  x  20  matrix
Generating  20  x  20  matrix
Generating  20  x  20  matrix
Generating  20  x  20  matrix
Generating  20  x  20  matrix
Generating  20  x  20  matrix
Generating  20  x  20  matrix
Generating  20  x  20  matrix
Generating  20  x  20  matrix
Generating  20  x  20  matrix
Generating  20  x  20  matrix
Generating  20  x  20  matrix
Generating  20  x  20  matrix
Generating  20  x  20  matrix
Generating  20  x  20  matrix
Generating  20  x  20  matrix
Generating  20  x  20  matrix
Generating  20  x  20  matrix
Generating  20  x  20  matrix
Generating  20  x  20  matrix
Generating  20  x  20  matrix
Generating  20  x  20  matrix
Generating  20  x  20  matrix
Generating  20  x  20  matrix
Generating  20  x  20  matrix
Generating  20  x  20  matrix
Generating  20  x  20  matrix
Generating  20  x  20  matrix
Generating  20  x  20  matrix
Generating  20  x  20  matrix
Generating  20  x  20  matrix
Generating  20  x  20  matrix
Generating  20  x  20  matrix
Generating  20  x  20  matrix
Generating  20  x  20  matrix
Generating  20  x  20  matrix
Generating  20  x  20  matrix
Generating  20  x  20  matrix
Generating  20  x  20  matrix
Generating  20  x  20  matrix
Generating  20  x  20  matrix
Generating  20  x  20  matrix
Generating  20  x  20  matrix
Generating  20  x  20  matrix
Generating  20  x  20  matrix
Generating  20  x  20  matrix
Generating  20  x  20  matrix
Generating  20  x  20  matrix
Generating  20  x  20  matrix
Generating  20  x  20  matrix
Generating  20  x  20  matrix
Generating  20  x  20  matrix
Generating  20  x  20  matrix
Generating  20  x  20  matrix
Generating  20  x  20  matrix
Generating  20  x  20  matrix
Generating  20  x  20  matrix
Generating  20  x  20  matrix
Generating  20  x  20  matrix
Generating  20  x  20  matrix
Generating  20  x  20  matrix
Generating  20  x  20  matrix
Generating  20  x  20  matrix
Generating  20  x  20  matrix
Generating  20  x  20  matrix
Generating  20  x  20  matrix
Generating  20  x  20  matrix
Generating  20  x  20  matrix
Generating  20  x  20  matrix
Generating  20  x  20  matrix
Generating  20  x  20  matrix
Generating  20  x  20  matrix
Generating  20  x  20  matrix
Generating  20  x  20  matrix
Generating  20  x  20  matrix
Generating  20  x  20  matrix
Generating  20  x  20  matrix
Generating  20  x  20  matrix
Generating  20  x  20  matrix
Generating  20  x  20  matrix
Generating  20  x  20  matrix
Generating  20  x  20  matrix
Generating  20  x  20  matrix
Generating  20  x  20  matrix
Generating  20  x  20  matrix
Generating  20  x  20  matrix
Generating  20  x  20  matrix
Generating  20  x  20  matrix
Generating  20  x  20  matrix
Generating  20  x  20  matrix
Generating  20  x  20  matrix
Generating  20  x  20  matrix
Generating  20  x  20  matrix
Generating  20  x  20  matrix
Generating  20  x  20  matrix
Generating  20  x  20  matrix
Generating  20  x  20  matrix
Generating  20  x  20  matrix
Generating  20  x  20  matrix
Generating  20  x  20  matrix
Generating  20  x  20  matrix
Generating  20  x  20  matrix
Generating  20  x  20  matrix
Generating  20  x  20  matrix
Generating  20  x  20  matrix
Generating  20  x  20  matrix
Generating  20  x  20  matrix
Generating  20  x  20  matrix
Generating  20  x  20  matrix
Generating  20  x  20  matrix
Generating  20  x  20  matrix
Generating  20  x  20  matrix
Generating  20  x  20  matrix
Generating  20  x  20  matrix
Generating  20  x  20  matrix
Generating  20  x  20  matrix
Generating  20  x  20  matrix
Generating  20  x  20  matrix
Generating  20  x  20  matrix
Generating  20  x  20  matrix
Generating  20  x  20  matrix
Generating  20  x  20  matrix
Generating  20  x  20  matrix
Generating  20  x  20  matrix
Generating  20  x  20  matrix
Generating  20  x  20  matrix
Generating  20  x  20  matrix
Generating  20  x  20  matrix
Generating  20  x  20  matrix
Generating  20  x  20  matrix
Generating  20  x  20  matrix
Generating  20  x  20  matrix
Generating  20  x  20  matrix
Generating  20  x  20  matrix
Generating  20  x  20  matrix
Generating  20  x  20  matrix
Generating  20  x  20  matrix
Generating  20  x  20  matrix
Generating  20  x  20  matrix
Generating  20  x  20  matrix
Generating  20  x  20  matrix
Generating  20  x  20  matrix
Generating  20  x  20  matrix
Generating  20  x  20  matrix
Generating  20  x  20  matrix
Generating  20  x  20  matrix
Generating  20  x  20  matrix
Generating  20  x  20  matrix
Generating  20  x  20  matrix
Generating  20  x  20  matrix
Generating  20  x  20  matrix
Generating  20  x  20  matrix
Generating  20  x  20  matrix
Generating  20  x  20  matrix
Generating  20  x  20  matrix
Generating  20  x  20  matrix
Generating  20  x  20  matrix
Generating  20  x  20  matrix
Generating  20  x  20  matrix
Generating  20  x  20  matrix
Generating  20  x  20  matrix
Generating  20  x  20  matrix
Generating  20  x  20  matrix
Generating  20  x  20  matrix
Generating  20  x  20  matrix
Generating  20  x  20  matrix
Generating  20  x  20  matrix
Generating  20  x  20  matrix
Generating  20  x  20  matrix
Generating  20  x  20  matrix
Generating  20  x  20  matrix
Generating  20  x  20  matrix
Generating  20  x  20  matrix
Generating  20  x  20  matrix
Generating  20  x  20  matrix
Generating  20  x  20  matrix
Generating  20  x  20  matrix
Generating  20  x  20  matrix
Generating  20  x  20  matrix
Generating  20  x  20  matrix
Generating  20  x  20  matrix
Generating  20  x  20  matrix
Generating  20  x  20  matrix
Generating  20  x  20  matrix
Generating  20  x  20  matrix
Generating  20  x  20  matrix
Generating  20  x  20  matrix
Generating  20  x  20  matrix
Generating  20  x  20  matrix
Generating  20  x  20  matrix
Generating  20  x  20  matrix
Generating  20  x  20  matrix
Generating  20  x  20  matrix
Generating  20  x  20  matrix
Generating  20  x  20  matrix
Generating  20  x  20  matrix
Generating  20  x  20  matrix
Generating  20  x  20  matrix
Generating  20  x  20  matrix
Generating  20  x  20  matrix
Generating  20  x  20  matrix
Generating  20  x  20  matrix
Generating  20  x  20  matrix
Generating  20  x  20  matrix
Generating  20  x  20  matrix
Generating  20  x  20  matrix
Generating  20  x  20  matrix
Generating  20  x  20  matrix
Generating  20  x  20  matrix
Generating  20  x  20  matrix
Generating  20  x  20  matrix
Generating  20  x  20  matrix
Generating  20  x  20  matrix
Generating  20  x  20  matrix
Generating  20  x  20  matrix
Generating  20  x  20  matrix
Generating  20  x  20  matrix
Generating  20  x  20  matrix
Generating  20  x  20  matrix
Generating  20  x  20  matrix
Generating  20  x  20  matrix
Generating  20  x  20  matrix
Generating  20  x  20  matrix
Generating  20  x  20  matrix
Generating  20  x  20  matrix
Generating  20  x  20  matrix
Generating  20  x  20  matrix
Generating  20  x  20  matrix
Generating  20  x  20  matrix
Generating  20  x  20  matrix
Generating  20  x  20  matrix
Generating  20  x  20  matrix
Generating  20  x  20  matrix
Generating  20  x  20  matrix
Generating  20  x  20  matrix
Generating  20  x  20  matrix
Generating  20  x  20  matrix
Generating  20  x  20  matrix
Generating  20  x  20  matrix
Generating  20  x  20  matrix
Generating  20  x  20  matrix
Generating  20  x  20  matrix
Generating  20  x  20  matrix
Generating  20  x  20  matrix
Generating  20  x  20  matrix
Generating  20  x  20  matrix
Generating  20  x  20  matrix
Generating  20  x  20  matrix
Generating  20  x  20  matrix
Generating  20  x  20  matrix
Generating  20  x  20  matrix
Generating  20  x  20  matrix
Generating  20  x  20  matrix
Generating  20  x  20  matrix
Generating  20  x  20  matrix
Generating  20  x  20  matrix
Generating  20  x  20  matrix
Generating  20  x  20  matrix
Generating  20  x  20  matrix
Generating  20  x  20  matrix
Generating  20  x  20  matrix
Generating  20  x  20  matrix
Generating  20  x  20  matrix
Generating  20  x  20  matrix
Generating  20  x  20  matrix
Generating  20  x  20  matrix
Generating  20  x  20  matrix
Generating  20  x  20  matrix
Generating  20  x  20  matrix
Generating  20  x  20  matrix
Generating  20  x  20  matrix
Generating  20  x  20  matrix
Generating  20  x  20  matrix
Generating  20  x  20  matrix
Generating  20  x  20  matrix
Generating  20  x  20  matrix
Generating  20  x  20  matrix
Generating  20  x  20  matrix
Generating  20  x  20  matrix
Generating  20  x  20  matrix
Generating  20  x  20  matrix
Generating  20  x  20  matrix
Generating  20  x  20  matrix
Generating  20  x  20  matrix
Generating  20  x  20  matrix
Generating  20  x  20  matrix
Generating  20  x  20  matrix
Generating  20  x  20  matrix
Generating  20  x  20  matrix
Generating  20  x  20  matrix
Generating  20  x  20  matrix
Generating  20  x  20  matrix
Generating  20  x  20  matrix
Generating  20  x  20  matrix
Generating  20  x  20  matrix
Generating  20  x  20  matrix
Generating  20  x  20  matrix
Generating  20  x  20  matrix
Generating  20  x  20  matrix
Generating  20  x  20  matrix
Generating  20  x  20  matrix
Generating  20  x  20  matrix
Generating  20  x  20  matrix
Generating  20  x  20  matrix
Generating  20  x  20  matrix
Generating  20  x  20  matrix
Generating  20  x  20  matrix
Generating  20  x  20  matrix
Generating  20  x  20  matrix
Generating  20  x  20  matrix
Generating  20  x  20  matrix
Generating  20  x  20  matrix
Generating  20  x  20  matrix
Generating  20  x  20  matrix
Generating  20  x  20  matrix
Generating  20  x  20  matrix
Generating  20  x  20  matrix
Generating  20  x  20  matrix
Generating  20  x  20  matrix
Generating  20  x  20  matrix
Generating  20  x  20  matrix
Generating  20  x  20  matrix
Generating  20  x  20  matrix
Generating  20  x  20  matrix
Generating  20  x  20  matrix
Generating  20  x  20  matrix
Generating  20  x  20  matrix
Generating  20  x  20  matrix
Generating  20  x  20  matrix
Generating  20  x  20  matrix
Generating  20  x  20  matrix
Generating  20  x  20  matrix
Generating  20  x  20  matrix
Generating  20  x  20  matrix
Generating  20  x  20  matrix
Generating  20  x  20  matrix
Generating  20  x  20  matrix
Generating  20  x  20  matrix
Generating  20  x  20  matrix
Generating  20  x  20  matrix
Generating  20  x  20  matrix
Generating  20  x  20  matrix
Generating  20  x  20  matrix
Generating  20  x  20  matrix
Generating  20  x  20  matrix
Generating  20  x  20  matrix
Generating  20  x  20  matrix
Generating  20  x  20  matrix
Generating  20  x  20  matrix
Generating  20  x  20  matrix
Generating  20  x  20  matrix
Generating  20  x  20  matrix
Generating  20  x  20  matrix
Generating  20  x  20  matrix
Generating  20  x  20  matrix
Generating  20  x  20  matrix
Generating  20  x  20  matrix
Generating  20  x  20  matrix
Generating  20  x  20  matrix
Generating  20  x  20  matrix
Generating  20  x  20  matrix
Generating  20  x  20  matrix
Generating  20  x  20  matrix
Generating  20  x  20  matrix
Generating  20  x  20  matrix
Generating  20  x  20  matrix
Generating  20  x  20  matrix
Generating  20  x  20  matrix
Generating  20  x  20  matrix
Generating  20  x  20  matrix
Generating  20  x  20  matrix
Generating  20  x  20  matrix
Generating  20  x  20  matrix
Generating  20  x  20  matrix
Generating  20  x  20  matrix
Generating  20  x  20  matrix
Generating  20  x  20  matrix
Generating  20  x  20  matrix
Generating  20  x  20  matrix
Generating  20  x  20  matrix
Generating  20  x  20  matrix
Generating  20  x  20  matrix
Generating  20  x  20  matrix
Generating  20  x  20  matrix
Generating  20  x  20  matrix
Generating  20  x  20  matrix
Generating  20  x  20  matrix
Generating  20  x  20  matrix
Generating  20  x  20  matrix
Generating  20  x  20  matrix
Generating  20  x  20  matrix
Generating  20  x  20  matrix
Generating  20  x  20  matrix
Generating  20  x  20  matrix
Generating  20  x  20  matrix
Generating  20  x  20  matrix
Generating  20  x  20  matrix
Generating  20  x  20  matrix
Generating  20  x  20  matrix
Generating  20  x  20  matrix
Generating  20  x  20  matrix
Generating  20  x  20  matrix
Generating  20  x  20  matrix
Generating  20  x  20  matrix
Generating  20  x  20  matrix
Generating  20  x  20  matrix
Generating  20  x  20  matrix
Generating  20  x  20  matrix
Generating  20  x  20  matrix
Generating  20  x  20  matrix
Generating  20  x  20  matrix
Generating  20  x  20  matrix
Generating  20  x  20  matrix
Generating  20  x  20  matrix
Generating  20  x  20  matrix
Generating  20  x  20  matrix
Generating  20  x  20  matrix
Generating  20  x  20  matrix
Generating  20  x  20  matrix
Generating  20  x  20  matrix
Generating  20  x  20  matrix
Generating  20  x  20  matrix
Generating  20  x  20  matrix
Generating  20  x  20  matrix
Generating  20  x  20  matrix
Generating  20  x  20  matrix
Generating  20  x  20  matrix
Generating  20  x  20  matrix
Generating  20  x  20  matrix
Generating  20  x  20  matrix
Generating  20  x  20  matrix
Generating  20  x  20  matrix
Generating  20  x  20  matrix
Generating  20  x  20  matrix
Generating  20  x  20  matrix
Generating  20  x  20  matrix
Generating  20  x  20  matrix
Generating  20  x  20  matrix
Generating  20  x  20  matrix
Generating  20  x  20  matrix
Generating  20  x  20  matrix
Generating  20  x  20  matrix
Generating  20  x  20  matrix
Generating  20  x  20  matrix
Generating  20  x  20  matrix
Generating  20  x  20  matrix
Generating  20  x  20  matrix
Generating  20  x  20  matrix
Generating  20  x  20  matrix
Generating  20  x  20  matrix
Generating  20  x  20  matrix
Generating  20  x  20  matrix
Generating  20  x  20  matrix
Generating  20  x  20  matrix
Generating  20  x  20  matrix
Generating  20  x  20  matrix
Generating  20  x  20  matrix
Generating  20  x  20  matrix
Generating  20  x  20  matrix
Generating  20  x  20  matrix
Generating  20  x  20  matrix
Generating  20  x  20  matrix
Generating  20  x  20  matrix
Generating  20  x  20  matrix
Generating  20  x  20  matrix
Generating  20  x  20  matrix
Generating  20  x  20  matrix
Generating  20  x  20  matrix
Generating  20  x  20  matrix
Generating  20  x  20  matrix
Generating  20  x  20  matrix
Generating  20  x  20  matrix
Generating  20  x  20  matrix
Generating  20  x  20  matrix
Generating  20  x  20  matrix
Generating  20  x  20  matrix
Generating  20  x  20  matrix
Generating  20  x  20  matrix
Generating  20  x  20  matrix
Generating  20  x  20  matrix
Generating  20  x  20  matrix
Generating  20  x  20  matrix
Generating  20  x  20  matrix
Generating  20  x  20  matrix
Generating  20  x  20  matrix
Generating  20  x  20  matrix
Generating  20  x  20  matrix
Generating  20  x  20  matrix
Generating  20  x  20  matrix
Generating  20  x  20  matrix
Generating  20  x  20  matrix
Generating  20  x  20  matrix
Generating  20  x  20  matrix
Generating  20  x  20  matrix
Generating  20  x  20  matrix
Generating  20  x  20  matrix
Generating  20  x  20  matrix
Generating  20  x  20  matrix
Generating  20  x  20  matrix
Generating  20  x  20  matrix
Generating  20  x  20  matrix
Generating  20  x  20  matrix
Generating  20  x  20  matrix
Generating  20  x  20  matrix
Generating  20  x  20  matrix
Generating  20  x  20  matrix
Generating  20  x  20  matrix
Generating  20  x  20  matrix
Generating  20  x  20  matrix
Generating  20  x  20  matrix
Generating  20  x  20  matrix
Generating  20  x  20  matrix
Generating  20  x  20  matrix
Generating  20  x  20  matrix
Generating  20  x  20  matrix
Generating  20  x  20  matrix
Generating  20  x  20  matrix
Generating  20  x  20  matrix
Generating  20  x  20  matrix
Generating  20  x  20  matrix
Generating  20  x  20  matrix
Generating  20  x  20  matrix
Generating  20  x  20  matrix
Generating  20  x  20  matrix
Generating  20  x  20  matrix
Generating  20  x  20  matrix
Generating  20  x  20  matrix
Generating  20  x  20  matrix
Generating  20  x  20  matrix
Generating  20  x  20  matrix
```


```
write.csv(results, "results/results.nodal.csv")
results
```


```
results.nodal <- read.csv("results/results.nodal.csv")

results.nodal$method <- factor(results.nodal$method, levels=c("OLS", "Node-label Permutation"))

p1 <- ggplot(results.nodal, aes(x=method, y=p_value,  fill=as.factor(effect))) +
  geom_violin(scale="width") +
  scale_fill_manual(values = c(cp[3], cp[8])) +
  geom_abline(slope=0, intercept=0.05, linetype="dashed") +
  labs(x="Method", y="p-value", fill="Effect", title="Nodal regression - Trait-based sex differences") +
  theme_classic()

p1

ggsave("figures/nodal.png", width=8, height=6)
```


```
aggregate(results.nodal$p_value, function(x) mean(x < 0.05), by=list(results.nodal$method, results.nodal$effect))
```


```
agg <- aggregate(p_value ~ method + effect, results.nodal, quantile)
do.call(data.frame, agg)
```


```
pvals_lm <- rep(0, 1)
pvals_perm <- rep(0, 1)

results <- data.frame(p_value=numeric(), method=numeric(), effect=numeric())

# Set parameters
areas <- 1
N <- 20
sampling_periods <- 100

for (effect in c(TRUE, FALSE)) {
  for (iter in 1:100) {
    # Generate nodes
    ids <- data.frame(ID=1:(areas*N),AREA=rep(1:areas,each=N),DEG_DIST=NA)
    
    # Generate a degree distribution and normalise it
    for (i in 1:areas) {
    degree_distribution <- rpois(N,3)+i
    ids$DEG_DIST[ids$AREA==i] <- degree_distribution/max(degree_distribution)
    }
    
    # Generate attributes
    # This creates a correlation between sex and underlying degree probability
    if (effect) {
      ids$SEX <- sapply(ids$DEG_DIST,FUN=function(x) { sample(c("M","F"),1,prob=c(x,1-x))})
    } else {
      ids$SEX <- sample(c("M","F"),N*areas,replace=TRUE)
    }
    # Generate probability for each edge (only within area)
    probs <- outer(ids$DEG_DIST, ids$DEG_DIST, "*")*outer(ids$AREA,ids$AREA,"==")
    
    # Create sampling periods, each sample only contains data from one area
    sps <- array(0,c(areas*sampling_periods,N*areas,N*areas))
    
    for (i in 1:areas) {
    sps[((sampling_periods*(i-1))+1):(sampling_periods*i),
    ((N*(i-1))+1):(N*i),((N*(i-1))+1):(N*i)] <- rgraph(N,m=sampling_periods,tprob=probs[((N*(i-1))+1):(N*i),
    ((N*(i-1))+1):(N*i)],mode="graph")
    }
    
    # Generate network
    network <- get_network(sps,data_format="SP")
    rownames(network) <- ids$SEX
    colnames(network) <- ids$SEX
    
    # Calculate degrees
    ids$DEGREE <- degree(network,gmode="graph")
    
    pval_lm <- summary(lm(DEGREE ~ SEX, data=ids))$coefficients[2, 4]
    pval_perm <- node_permutation(ids$DEGREE, as.numeric(factor(ids$SEX)) - 1)
    
    results[nrow(results) + 1, ] <- list(pval_lm, "OLS", as.character(effect))
    results[nrow(results) + 1, ] <- list(pval_perm, "Node-label Permutation", as.character(effect))
  }
}

write.csv(results, "results/results.nodal.effect.csv")
results
```
