## Supplementary material for "Common Permutation Methods in Animal Social Network Analysis Do Not Control for Non-independence": Simulation code: nodal_regression_dependence.nb.html

R Notebook


Code 

- Show All Code
- Hide All Code
- Download Rmd

### R Notebook


```
node_permutation <- function(y, x) {
  obs_b <- lm(y ~ x)$coefficients[2]
  null_b <- sapply(1:10000, function(i) .lm.fit(cbind(1, x), sample(y))$coefficients[2])
  mean(abs(obs_b) < abs(null_b))
}

weighted_sbm <- function(n) {
  M <- sample(1:2, n, replace=TRUE, prob=c(0.5, 0.5))
  A <- matrix(0, n, n)
  for (i in 1:n) {
    for (j in 1:n) {
      if (i < j) {
        if (M[i] == M[j]) {
          A[i, j] <- rbinom(1, 1, 0.8) * runif(1, min=0)
        } else {
          A[i, j] <- rbinom(1, 1, 0.4) * runif(1, max=1)
        }
      }
    }
  }
  list(A=A, M=M)
}
```


```
group_sizes <- c()

n <- 40
b <- 0.05

results <- data.frame(p_value=numeric(), method=numeric(), effect=numeric())
effect <- FALSE
for (effect in c(TRUE, FALSE)) {
  for (iter in 1:1000) {
    # Y_ <- matrix(rbinom(n^2, 1, 1.0) * runif(n^2), n, n)
    sbm_obj <- weighted_sbm(n)
    Y_ <- sbm_obj$A
    Y_ <- Y_ * upper.tri(Y_)
    Y_ <- Y_ + t(Y_)

    x <- runif(n) + runif(2)[sbm_obj$M]
    y <- rowSums(Y_)
    
    group_sizes <- c(group_sizes, sum(sbm_obj$M == 1))
    
    if (effect) {
      x <- b * y + (1 - b) * x
    }

    pval_lm <- summary(lm(y ~ x))$coefficients[2, 4]
    pval_perm <- node_permutation(y, x)
    
    results[nrow(results) + 1, ] <- list(pval_lm, "OLS", as.character(effect))
    results[nrow(results) + 1, ] <- list(pval_perm, "Node-label Permutation", as.character(effect))
  }
}

write.csv(results, "results/results.nodal.clique.csv")
results
```


```
results.nodal.clique <- read.csv("results/results.nodal.clique.csv")

ggplot(results.nodal.clique, aes(x=as.factor(method), y=p_value,  fill=as.factor(effect))) +
  geom_violin(scale="width") +
  scale_fill_manual(values = c(cp[3], cp[8])) +
  geom_abline(slope=0, intercept=0.05, linetype="dashed") +
  labs(x="Method", y="p-value", fill="Effect", title = "Nodal regression - Dependence on substructure") +
  theme_classic()
```


### Group membership cannot be reliably detected


```
sbm_obj <- weighted_sbm(20)
G <- graph_from_adjacency_matrix(sbm_obj$A, mode="undirected", weighted=TRUE)

membership(cluster_fast_greedy(G))[sbm_obj$M == 1]
```


```
 [1] 1 2 2 1 1 1 3 2 2 1 1
```


```
membership(cluster_fast_greedy(G))[sbm_obj$M == 2]
```


```
[1] 3 1 3 2 2 1 2 3 3
```
