## Supplementary material for "Common Permutation Methods in Animal Social Network Analysis Do Not Control for Non-independence": Simulation code: plot_figures.nb.html


Code 

- Show All Code
- Hide All Code
- Download Rmd

### Plot figures


```
library(ggplot2)
library(patchwork)
```


```
results.nodal <- read.csv("results/results.nodal.csv")
results.dyadic <- read.csv("results/results.dyadic.csv")
results.nodal.dependence <- read.csv("results/results.nodal.dependence.csv")
results.dyadic.dependence <- read.csv("results/results.dyadic.dependence.csv")
```


```
results <- read.csv("results/results.nodal.csv")

results$effect <- as.factor(results$effect)
levels(results$effect) <- c("No", "Yes")
levels(results$method) <- c("Node-label Permutation", "LM")

results$effect <- factor(results$effect, levels=c("Yes", "No"))
results$method <- factor(results$method, levels=c("LM", "Node-label Permutation"))

df_text <- aggregate(cbind(power=p_value) ~ method + effect, results, function(x) mean(x < 0.05))
df_text$label <- ""
for (i in 1:nrow(df_text)) {
  if (df_text[i, ]$effect == "Yes") {
    df_text[i, ]$label <- paste0("True Positives: ", signif(df_text[i, ]$power * 100, 3), "%")
  } else {
    df_text[i, ]$label <- paste0("False Positives: ", signif(df_text[i, ]$power * 100, 3), "%")
  }
}

p1 <- ggplot(results, aes(x=p_value, fill=effect)) +
  geom_density() +
  scale_fill_manual(values = c(cp[6], cp[8])) +
  facet_grid(rows=vars(method), cols=vars(effect), scales="free") +
  geom_vline(xintercept=0.05, linetype="dashed") +
  labs(x = "p-value", y="Density", fill="Effect", title = "Nodal - Strength Differences") +
  geom_text(
    data    = df_text,
    mapping = aes(x = Inf, y = Inf, label = label),
    hjust   = 1.05,
    vjust   = 1.5
  ) +
  xlim(0, 1) +
  theme_classic() +
  theme(strip.background = element_rect(colour="white", fill="#f5f5f5"), strip.text.x = element_blank())

p1
```


```
results <- read.csv("results/results.nodal.dependence.csv")

results$effect <- as.factor(results$effect)
levels(results$effect) <- c("No", "Yes")
levels(results$method) <- c("Node-label Permutation", "LM")

results$effect <- factor(results$effect, levels=c("Yes", "No"))
results$method <- factor(results$method, levels=c("LM", "Node-label Permutation"))

df_text <- aggregate(cbind(power=p_value) ~ method + effect, results, function(x) mean(x < 0.05))
df_text$label <- ""
for (i in 1:nrow(df_text)) {
  if (df_text[i, ]$effect == "Yes") {
    df_text[i, ]$label <- paste0("True Positives: ", signif(df_text[i, ]$power * 100, 3), "%")
  } else {
    df_text[i, ]$label <- paste0("False Positives: ", signif(df_text[i, ]$power * 100, 3), "%")
  }
}

p2 <- ggplot(results, aes(x=p_value, fill=effect)) +
  geom_density() +
  scale_fill_manual(values = c(cp[6], cp[8])) +
  facet_grid(rows=vars(method), cols=vars(effect), scales="free") +
  geom_vline(xintercept=0.05, linetype="dashed") +
  labs(x = "p-value", y="Density", fill="Effect", title = "Nodal - Clique Dependence") +
  geom_text(
    data    = df_text,
    mapping = aes(x = Inf, y = Inf, label = label),
    hjust   = 1.05,
    vjust   = 1.5
  ) +
  xlim(0, 1) +
  theme_classic() +
  theme(strip.background = element_rect(colour="white", fill="#f5f5f5"), strip.text.x = element_blank())

p2
```


```
results <- read.csv("results/results.dyadic.csv")

results$effect <- as.factor(results$effect)
levels(results$effect) <- c("No", "Yes")
levels(results$method) <- c("LM", "MMLM", "QAP")

results$effect <- factor(results$effect, levels=c("Yes", "No"))
results$method <- factor(results$method, levels=c("LM", "QAP", "MMLM"))

df_text <- aggregate(cbind(power=p_value) ~ method + effect, results, function(x) mean(x < 0.05))
df_text$label <- ""
for (i in 1:nrow(df_text)) {
  if (df_text[i, ]$effect == "Yes") {
    df_text[i, ]$label <- paste0("True Positives: ", signif(df_text[i, ]$power * 100, 3), "%")
  } else {
    df_text[i, ]$label <- paste0("False Positives: ", signif(df_text[i, ]$power * 100, 3), "%")
  }
}

p3 <- ggplot(results, aes(x=p_value, fill=effect)) +
  geom_density() +
  scale_fill_manual(values = c(cp[6], cp[8])) +
  facet_grid(rows=vars(method), cols=vars(effect), scales="free") +
  geom_vline(xintercept=0.05, linetype="dashed") +
  labs(x = "p-value", y="Density", fill="Effect", title = "Dyadic - Node Dependence") +
  geom_text(
    data    = df_text,
    mapping = aes(x = Inf, y = Inf, label = label),
    hjust   = 1.05,
    vjust   = 1.5
  ) +
  xlim(0, 1) +
  theme_classic() +
  theme(strip.background = element_rect(colour="white", fill="#f5f5f5"), strip.text.x = element_blank())

p3
```


```
results <- read.csv("results/results.dyadic.dependence.csv")

results$effect <- as.factor(results$effect)
levels(results$effect) <- c("No", "Yes")
levels(results$method) <- c("LM", "MMLM", "QAP")

results$effect <- factor(results$effect, levels=c("Yes", "No"))
results$method <- factor(results$method, levels=c("LM", "QAP", "MMLM"))

df_text <- aggregate(cbind(power=p_value) ~ method + effect, results, function(x) mean(x < 0.05))
df_text$label <- ""
for (i in 1:nrow(df_text)) {
  if (df_text[i, ]$effect == "Yes") {
    df_text[i, ]$label <- paste0("True Positives: ", signif(df_text[i, ]$power * 100, 3), "%")
  } else {
    df_text[i, ]$label <- paste0("False Positives: ", signif(df_text[i, ]$power * 100, 3), "%")
  }
}

p4 <- ggplot(results, aes(x=p_value, fill=effect)) +
  geom_density() +
  scale_fill_manual(values = c(cp[6], cp[8])) +
  facet_grid(rows=vars(method), cols=vars(effect), scales="free") +
  geom_vline(xintercept=0.05, linetype="dashed") +
  labs(x = "p-value", y="Density", fill="Effect", title = "Dyadic - Clique Dependence") +
  geom_text(
    data    = df_text,
    mapping = aes(x = Inf, y = Inf, label = label),
    hjust   = 1.05,
    vjust   = 1.5
  ) +
  xlim(0, 1) +
  theme_classic() +
  theme(strip.background = element_rect(colour="white", fill="#f5f5f5"), strip.text.x = element_blank())

p4
```


```
p1 + p2 + p3 + p4 + plot_layout(guides="collect") & theme(legend.position = 'bottom')
ggsave("figures/combined.pvalue.png", width=10, height=8, dpi=700)
```


```
results <- read.csv("results/results.dyadic.csv")

results$effect <- as.factor(results$effect)
levels(results$effect) <- c("No", "Yes")
levels(results$method) <- c("LM", "MMLM", "QAP")

results$effect <- factor(results$effect, levels=c("Yes", "No"))
results$method <- factor(results$method, levels=c("LM", "QAP", "MMLM"))

e1 <- ggplot(results, aes(x=effect_size, fill=effect)) +
  geom_density() +
  scale_fill_manual(values = c(cp[6], cp[8])) +
  facet_grid(rows=vars(method), cols=vars(effect), scales="free") +
  geom_vline(xintercept=0.0, linetype="dashed") +
  labs(x = "Effect Size Estimate", y="Density", fill="Effect", title = "Node Dependence") +
  theme_classic() +
  theme(strip.background = element_rect(colour="white", fill="#f5f5f5"), strip.text.x = element_blank())
```


```
results <- read.csv("results/results.dyadic.dependence.csv")

results$effect <- as.factor(results$effect)
levels(results$effect) <- c("No", "Yes")
levels(results$method) <- c("LM", "MMLM", "QAP")

results$effect <- factor(results$effect, levels=c("Yes", "No"))
results$method <- factor(results$method, levels=c("LM", "QAP", "MMLM"))

e2 <- ggplot(results, aes(x=effect_size, fill=effect)) +
  geom_density() +
  scale_fill_manual(values = c(cp[6], cp[8])) +
  facet_grid(rows=vars(method), cols=vars(effect), scales="free") +
  geom_vline(xintercept=0.0, linetype="dashed") +
  labs(x = "Effect Size Estimate", y="Density", fill="Effect", title = "Clique Dependence") +
  theme_classic() +
  theme(strip.background = element_rect(colour="white", fill="#f5f5f5"), strip.text.x = element_blank())
```


```
e1 + e2 + plot_layout(guides="collect") & theme(legend.position = 'bottom')
ggsave("figures/combined.effect.png", width=12, height=8, dpi=700)
```


```
lambda <- seq(0.1, 10, 0.1)
x <- 2; d <- 1
probs_1 <- dpois(x, lambda * d)
x <- 10; d <- 5
probs_2 <- dpois(x, lambda * d)

qplot(lambda, probs_1, geom="line") +
  geom_area(fill=cp[2], alpha=0.4, color="black") +
  geom_area(aes(x=lambda, y=probs_2/max(probs_1)), fill=cp[1], alpha=0.4, color="black") +
  geom_vline(xintercept=2, linetype="dashed") +
  labs(x="Interaction rate", y="Probability (normalised)") +
  geom_text(aes(x=3.5, y=0.35), label="X = 10, D = 5") +
  geom_text(aes(x=6, y=0.1), label="X = 2, D = 1") +
  theme_classic()
```


```
# quantile(sample(lambda, 1e6, replace=TRUE, prob=probs), probs=c(0.025, 0.50, 0.975))
```


```
x <- 2
d <- 1

lambda <- seq(0.01, 5, 0.01)
probs <- dpois(x, lambda * d)
plot(lambda, probs, type="l")
```


```
quantile(sample(lambda, 1e6, replace=TRUE, prob=probs), probs=c(0.025, 0.50, 0.975))
```


```
 2.5%   50% 97.5% 
 0.59  2.43  4.76
```


```
p1 + p2 + p3 + p4 + plot_layout(guides="collect") & theme(legend.position = 'bottom')
ggsave("figures/combined.pvalue.png", width=10, height=8, dpi=700)
```


```
results <- read.csv("results/results.dyadic.csv")

results$effect <- as.factor(results$effect)
levels(results$effect) <- c("No", "Yes")
levels(results$method) <- c("LM", "MMLM", "QAP")

results$effect <- factor(results$effect, levels=c("Yes", "No"))
results$method <- factor(results$method, levels=c("LM", "QAP", "MMLM"))

e1 <- ggplot(results, aes(x=effect_size, fill=effect)) +
  geom_density() +
  scale_fill_manual(values = c(cp[6], cp[8])) +
  facet_grid(rows=vars(method), cols=vars(effect), scales="free") +
  geom_vline(xintercept=0.0, linetype="dashed") +
  labs(x = "Effect Size Estimate", y="Density", fill="Effect", title = "Node Dependence") +
  theme_classic() +
  theme(strip.background = element_rect(colour="white", fill="#f5f5f5"), strip.text.x = element_blank())
```


```
results <- read.csv("results/results.dyadic.dependence.csv")

results$effect <- as.factor(results$effect)
levels(results$effect) <- c("No", "Yes")
levels(results$method) <- c("LM", "MMLM", "QAP")

results$effect <- factor(results$effect, levels=c("Yes", "No"))
results$method <- factor(results$method, levels=c("LM", "QAP", "MMLM"))

e2 <- ggplot(results, aes(x=effect_size, fill=effect)) +
  geom_density() +
  scale_fill_manual(values = c(cp[6], cp[8])) +
  facet_grid(rows=vars(method), cols=vars(effect), scales="free") +
  geom_vline(xintercept=0.0, linetype="dashed") +
  labs(x = "Effect Size Estimate", y="Density", fill="Effect", title = "Clique Dependence") +
  theme_classic() +
  theme(strip.background = element_rect(colour="white", fill="#f5f5f5"), strip.text.x = element_blank())
```


```
e1 + e2 + plot_layout(guides="collect") & theme(legend.position = 'bottom')
ggsave("figures/combined.effect.png", width=10, height=5, dpi=700)
```


```
lambda <- seq(0.1, 10, 0.1)
x <- 2; d <- 1
probs_1 <- dpois(x, lambda * d)
x <- 10; d <- 5
probs_2 <- dpois(x, lambda * d)

qplot(lambda, probs_1, geom="line") +
  geom_area(fill=cp[2], alpha=0.4, color="black") +
  geom_area(aes(x=lambda, y=probs_2/max(probs_1)), fill=cp[1], alpha=0.4, color="black") +
  geom_vline(xintercept=2, linetype="dashed") +
  labs(x="Interaction rate", y="Probability (normalised)") +
  geom_text(aes(x=3.5, y=0.35), label="X = 10, D = 5") +
  geom_text(aes(x=6, y=0.1), label="X = 2, D = 1") +
  theme_classic()

# quantile(sample(lambda, 1e6, replace=TRUE, prob=probs), probs=c(0.025, 0.50, 0.975))
```


```
x <- 2
d <- 1

lambda <- seq(0.01, 5, 0.01)
probs <- dpois(x, lambda * d)
plot(lambda, probs, type="l")

quantile(sample(lambda, 1e6, replace=TRUE, prob=probs), probs=c(0.025, 0.50, 0.975))
```


```
lk.pois <- function(par, x) {
  -dpois(x, par)
}
```
