## Supplementary figures and images for "Common Permutation Methods in Animal Social Network Analysis Do Not Control for Non-independence"

### combined.effect.png

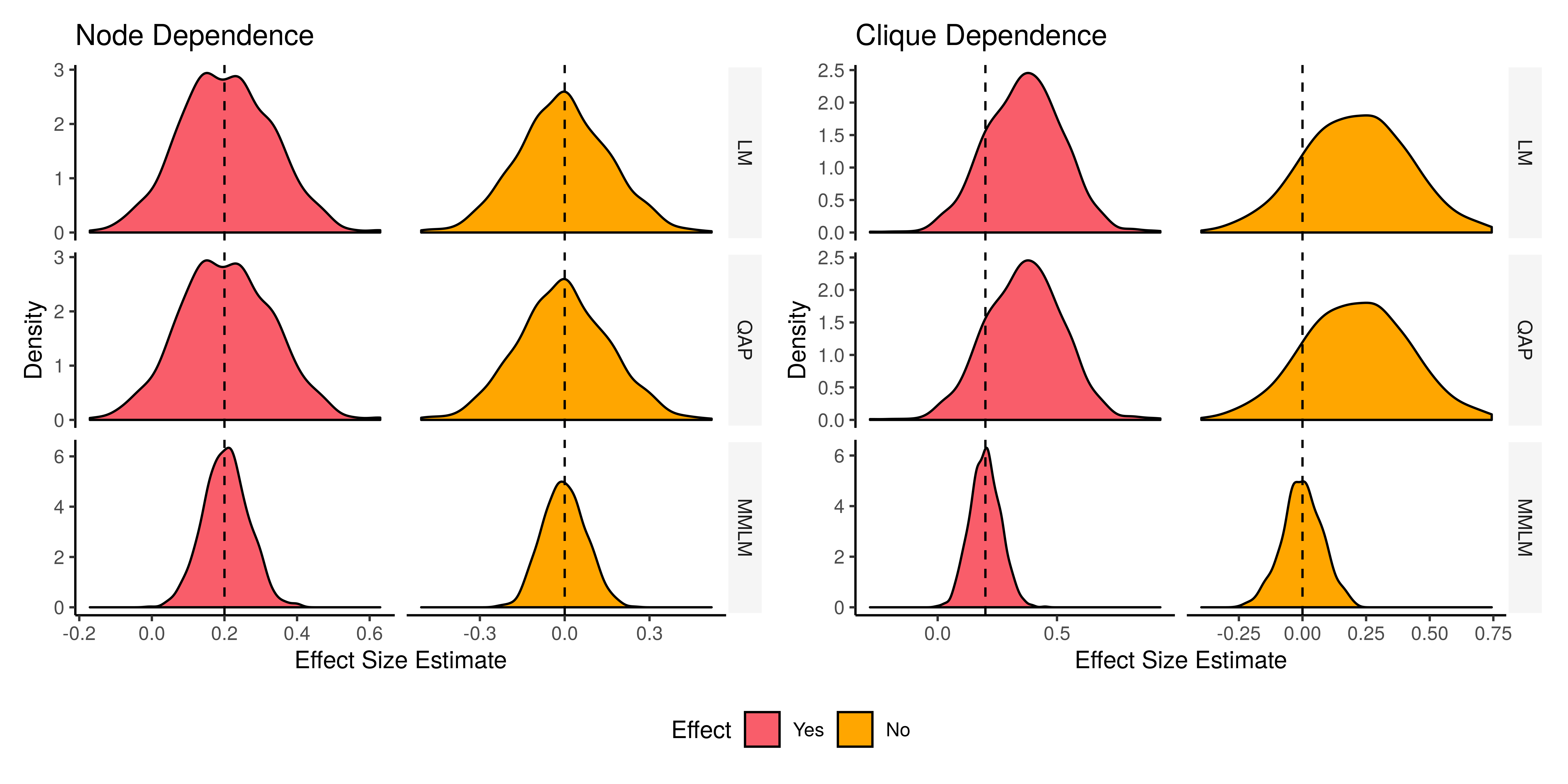

### combined.pvalue.png

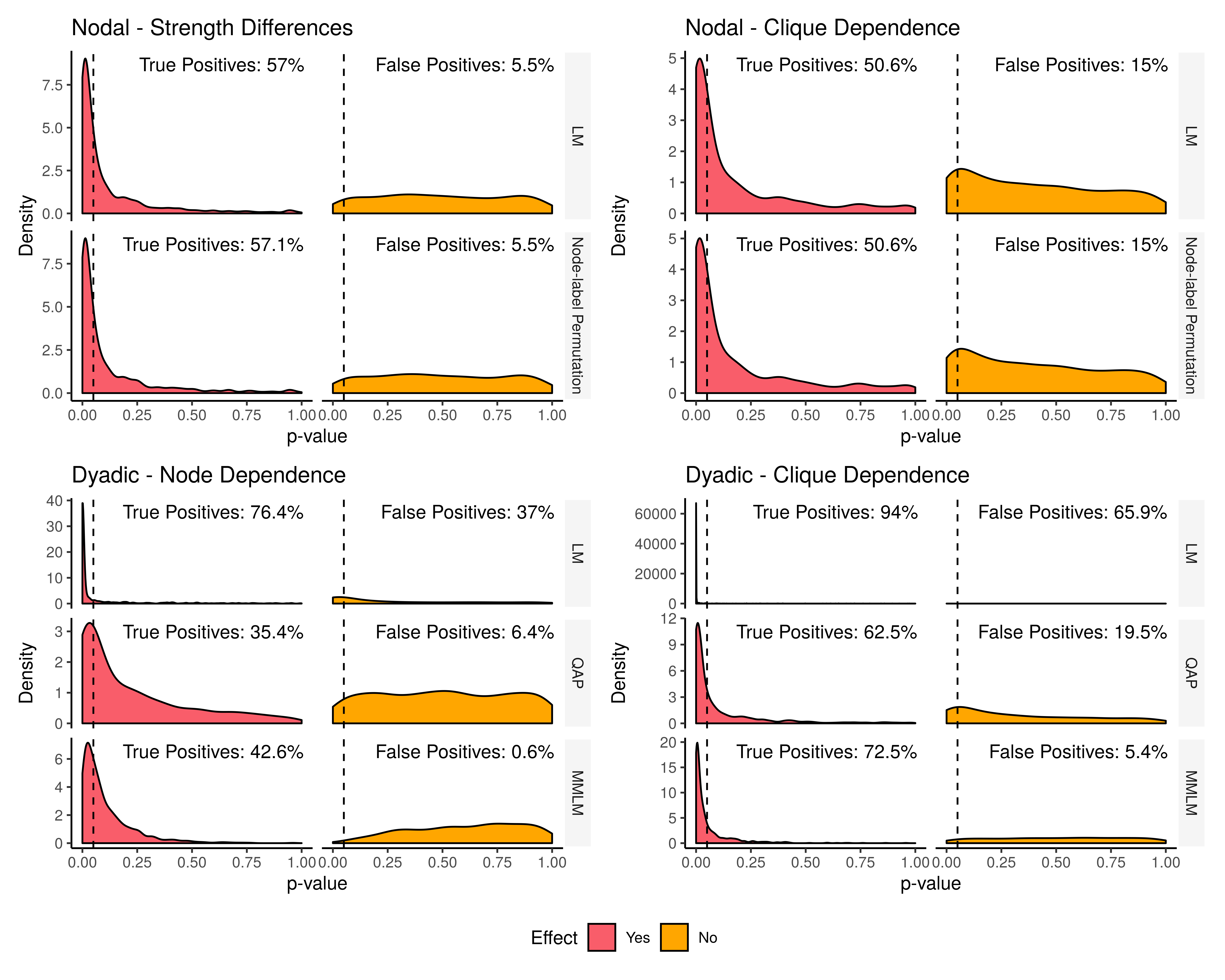
